# Osmotic stress response of the coral and oyster pathogen *Vibrio coralliilyticus*: acquisition of catabolism gene clusters for the compatible solute and signaling molecule *myo* -inositol

**DOI:** 10.1101/2024.01.16.575920

**Authors:** Katherine E. Boas Lichty, Rachel M. Loughran, Blake Ushijima, Gary P. Richards, E. Fidelma Boyd

## Abstract

Marine bacteria experience fluctuations in osmolarity that they must adapt to, and most bacteria respond to high osmolarity by accumulating compatible solutes also known as osmolytes. The osmotic stress response and compatible solutes used by the coral and oyster pathogen *Vibrio coralliilyticus* were unknown. In this study, we showed that to alleviate osmotic stress *V. coralliilyticus* biosynthesized glycine betaine (GB) and transported into the cell choline, GB, ectoine, dimethylglycine, and dimethylsulfoniopropionate, but not *myo*-inositol. *Myo*-inositol is a stress protectant and a signaling molecule that is biosynthesized and used by algae. Bioinformatics identified *myo*-inositol (*iol*) catabolism clusters in *V. coralliilyticus* and other *Vibrio, Photobacterium, Grimontia,* and *Enterovibrio* species. Growth pattern analysis demonstrated that *V. coralliilyticus* utilized *myo*-inositol as a sole carbon source, with a short lag time of 3 h. An *iolG* deletion mutant, which encodes an inositol dehydrogenase, was unable to grow on *myo*-inositol. Within the *iol* clusters were an MFS-type (*iolT1)* and an ABC-type (*iolXYZ)* transporter and analyses showed that both transported *myo*-inositol. IolG and IolA phylogeny among *Vibrionaceae* species showed different evolutionary histories indicating multiple acquisition events. Outside of *Vibrionaceae*, IolG was most closely related to IolG from a small group of *Aeromonas* fish and human pathogens and *Providencia* species. However, IolG from hypervirulent *A. hydrophila* strains clustered with IolG from *Enterobacter,* and divergently from *Pectobacterium, Brenneria,* and *Dickeya* plant pathogens. The *iol* cluster was also present within *Aliiroseovarius, Burkholderia, Endozoicomonas, Halomonas, Labrenzia, Marinomonas, Marinobacterium, Cobetia, Pantoea,* and *Pseudomonas,* of which many species were associated with marine flora and fauna.

**IMPORTANCE:** Host associated bacteria such as *V. coralliilyticus* encounter competition for nutrients and have evolved metabolic strategies to better compete for food. Emerging studies show that *myo*-inositol is exchanged in the coral-algae symbiosis, is likely involved in signaling, but is also an osmolyte in algae. The bacterial consumption of *myo*-inositol could contribute to a breakdown of the coral-algae symbiosis during thermal stress or disrupt the coral microbiome. Phylogenetic analyses showed that the evolutionary history of *myo*-inositol metabolism is complex, acquired multiple times in *Vibrio,* but acquired once in many bacterial plant pathogens. Further analysis also showed that a conserved *iol* cluster is prevalent among many marine species (commensals, mutualists, and pathogens) associated with marine flora and fauna, algae, sponges, corals, molluscs, crustaceans, and fish.

## INTRODUCTION

*Vibrio coralliilyticus* is a marine bacterium, a member of the family *Vibrionaceae*, an important pathogen of coral, and along with *V. mediterranei (*previously *V. shilonii)* a source of coral disease worldwide (1–10). This species was first characterized as an etiological agent for coral bleaching and tissue loss of the coral *Pocillopora damicornis*, and has since been identified as a pathogen in numerous other species of coral (2, 3, 6, 11–23). Recent studies also showed that *V. coralliilyticus* and a related species *V. tubiashii* caused high mortalities in oyster larvae of both Pacific oysters (*Crassostrea gigas*) and Eastern oysters (*Crassostrea virginica)* in the United States (15, 24–29).

Vibriosis in corals as well as oyster larvae is linked to increased water temperatures, especially for select strains of the pathogen *V. coralliilyticus* (2, 23, 25). In response to elevated temperatures, *V. coralliilyticus* undergoes various changes to its transcriptome and proteome (11, 13), which includes various genes important for infections of hosts like corals (23). The increased temperatures also correlates with the increased abundance of *V. coralliilyticus* (5), and the structure of natural *Vibrio* populations could be altered within the coral holobiont (30–32). This restructuring is proposed to contribute to disruption of processes like the nutritional exchange within the coral (33) as well as the healthy coral microbiome (34, 35). One consequence of infection is bleaching, the loss of the corals’ endosymbiotic dinoflagellates (family *Symbiodiniaceae*), that occurs with select pathogenic strains of *V. coralliilyticus* (2, 17). The photosynthetic dinoflagellate cells participate in a nutrient exchange with the host coral cells, which serves as the main energy source for many reef-building coral species (36–41). This metabolic exchange with photosynthetic dinoflagellates is central to maintaining the health of this mutualistic relationship (42–47).

One metabolite exchanged between dinoflagellates and coral is the carbocyclic sugar *myo*-inositol hypothesized to be transported to coral with a function not yet established (37, 38, 48). *Myo*-inositol is a six-carbon cyclic polyol and the most abundant form of inositol in the environment. It is an important component of eukaryotic cell membranes as phosphatidylinositol (PI) and a signaling molecule (49–51). In dinoflagellates inositol isomers are biosynthesized as important compatible solutes and are significantly enriched during thermal stress (38, 48, 52–55). In response to increased salinity, algae, plants, and marine invertebrates use *myo*-inositol as a compatible solute for osmotic stress protection (56–58). Additionally, in algae and plants, *myo*-inositol has been implicated as an important signaling molecule (48, 54, 55, 59).

Studies have shown that soil bacteria and human bacterial pathogens can utilize inositols as carbon and energy sources (60, 61). The first step in the pathway requires inositol dehydrogenases (IDHs), which are specific for different isomers of inositol such as *myo*-, *scyllo*-, and d-*chiro*-inositol. Genetics and enzyme functionality studies for bacterial *myo*-inositol catabolism genes were first established and investigated in *Bacillus subtilis* (62–65) (**Fig. 1**). *Myo*-inositol is degraded by inositol 2-dehydrogenase (IolG) into 2-keto-*myo*-inositol, which is then converted by *myo*-inosose dehydratase (IolE) to 3,5/4-Trihydroxycylohexane-1,2-dione. This is subsequently hydrolyzed into 5- deoxy-D-gluconate by an acylhydrolase (IolD). This product is then isomerized by 5- deoxy-glucuronate isomerase (IolB) and phosphorylated by 5-dehydro-2- deoxygluconokinase (IolC). Then 2-deoxy-5-keto-D-gluconate-6-P is converted to dihydroxyacetone (DHA) and malonate semialdehyde by an aldolase (IolJ). Malonate semialdehyde is converted to acetyl-CoA by the CoA-acylating methylmalonate-semialdehyde dehydrogenase (IolA). The product DHA can enter glycolysis and acetyl-CoA can enter the tricarboxylic acid (TCA) cycle (63–66). The transport of *myo*-inositol into *B. subtilis* for catabolism is performed by two major facilitator superfamily (MFS)- type transporters, named IolT and IolF (63). Several Gram-negative bacteria have been shown to use *myo*-inositol as a metabolite with a similar pathway, species in the genera *Klebsiella, Caulobacter, Rhizobium, Sinorhizobium, Pseudomonas, Yersinia, Corynebacterium, Salmonella*, and *Serratia* (67–76). For example, the human pathogen *Salmonella enterica* serovar Typhimurium utilized *myo*-inositol as a sole carbon source, but growth on *myo*- inositol was characterized by a long lag phase of > 45 h and a bistable phenotype (61, 69, 70, 77, 78). This species uses two MFS family transporters, named IolT1 and IolT2 for *myo*-inositol uptake (70). In the Alpha-proteobacteria *Caulobacter vibrioides* (formerly *crescentus*) catabolism of *myo*-inositol was demonstrated and an ABC-type transporter was used for uptake (68, 79). *Myo*-inositol utilization has not been explored extensively among other marine bacteria.

**Fig. 1.**
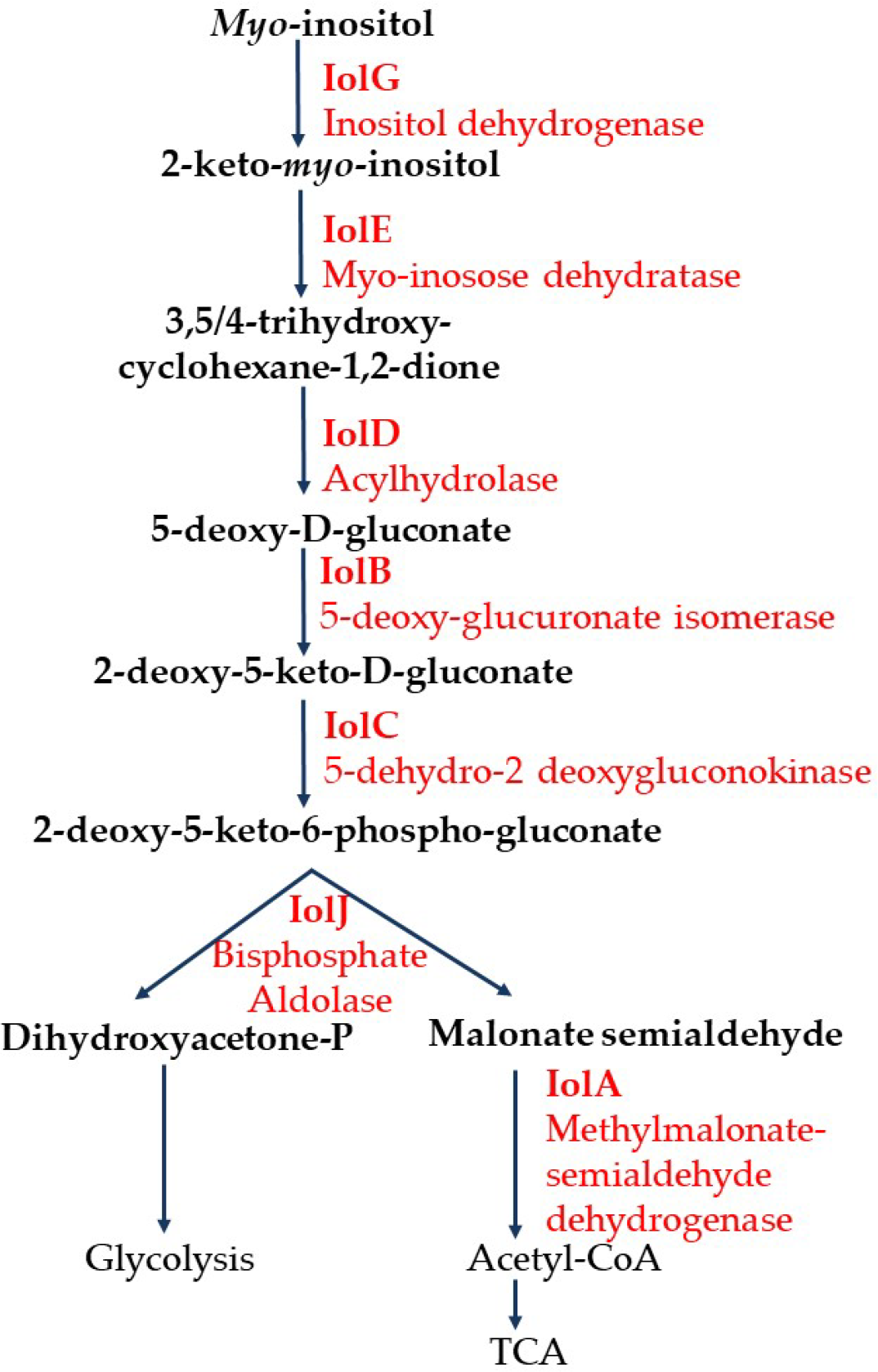
Pathway of m*yo*-inositol catabolism. The pathway presented is based upon studies in *B. subtilis* and *Salmonellla enterica*. Enzymes (red) and intermediate compounds in the pathway are displayed.

Here, we established osmotic stress response mechanisms using bioinformatics and genetic analyses among *V. coralliilyticus* strains CN52H-1 (16), OCN014 (23), BAA- 450 (19), and RE87 (80, 81) encompassing strains isolated from healthy and diseased hosts. We examined *V. coralliilyticus* growth patterns at different temperatures and salinities to determine their salinity tolerance. Bioinformatics and physiological approaches were employed to determine the compatible solutes used by of *V. coralliilyticus* for osmotic protection. Our investigations led to the identification of a large genomic region containing genes for *myo*-inositol catabolism, a molecule used in many eukaryotic cellular functions including stress protection and signaling. Using comparative genomics and phylogenetic analyses, the evolutionary history of *myo*- inositol catabolism genes among the *Vibrionaceae* was established. The distribution of *iol* clusters among marine bacteria was also examined.

## RESULTS

### *Vibrio coralliilyticus* salinity and temperature range

To determine the ability of *V. coralliilyticus* to grow in a range of NaCl concentrations and at different temperatures, growth patterns were examined in M9 minimal media supplemented with 20 mM glucose (M9G) and a final concentration of 1% to 6% NaCl at 28°C and 37°C for 24 h.

M9 minimal media was used for growth assays because it does not contain compatible solutes or their precursors and this allowed us to examine the response of *V. coralliilyticus* to increased salinity without the protection of external compatible solutes. These analyses showed that the *V. coralliilyticus* strains grew best in M9G 2% NaCl at 28°C (**Fig. 2**). For example, at this temperature, for strain CN52H-1, the final OD595 reading was 0.81 in 2% NaCl whereas in 1% NaCl, the final OD reached was 0.49. When the salinity was increased to 3% NaCl the maximum growth rate (0.62 h^-1^) was lower than what was observed at 2% NaCl (0.70 h^-1^). Growth of *V. coralliilyticus* in 4% NaCl showed a 2 h lag time and a lower maximum growth rate (0.33 h^-1^) whereas no growth occurred in 5% and 6% NaCl (**Fig. 2A**). Next, we tested whether increased temperature affected the salinity growth range by examining growth at 37°C (**Fig. 2B**). Interestingly, incubation at 37°C impacted the ability to grow in low salinity; at this temperature there was no growth in 1% NaCl (**Fig. 2B**). Growth pattern analysis was also performed on *V. coralliilyticus* OCN014 (**Fig. 2C-D**), BAA-450 (**Fig. 2E-F**), and RE87 (**Fig. 2G-H**), pathogenic strains of corals and oyster larvae in 1% to 6% NaCl at 28°C and 37°C for 24 h. We also examined the salinity growth range at 21°C and 34°C for strain CN52H-1 (**Fig. S1A-B)**. Two observations were made from these analyses; first incubation at 21°C negatively affected growth in 4% NaCl resulting in a longer lag phase than at any other temperature (**Fig. S1B)**. Secondly, growth at 34°C was similar to 28°C, but with a longer lag phase in M9G 4% NaCl (**Fig. S1A)**. These analyses demonstrated that *V. coralliilyticus* was unable to grow in M9G media with a salinity at or above 5% NaCl and at 37°C strains were unable to grow in 1% NaCl (**Fig. 2**). Overall, all strains showed similar osmotic stress tolerance responses.

**Fig. 2.**
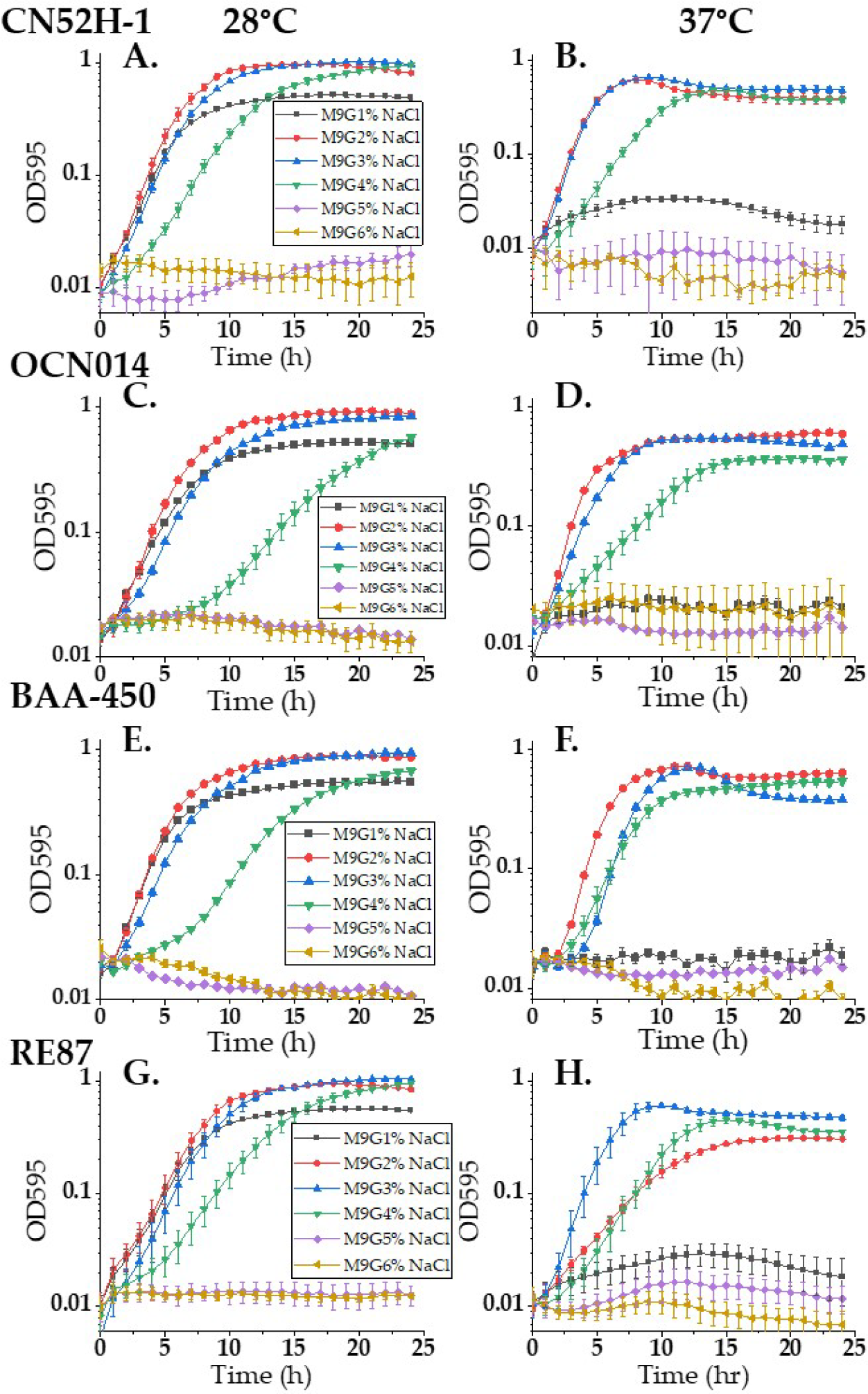
*Vibrio coralliilyticus* NaCl tolerance range. Growth curves of *V. coralliilyticus* strains in minimal media (M9) with glucose (M9G) with a final concentration of 1% to 6% NaCl. Growth was measured every hour for 24 h at **A.** 28°C strain CN52H-1, **B.** 37°C strain CN52H-1, **C.** 28°C strain OCN014, **D.** 37°C strain OCN014, **E.** 28°C strain BAA-450, **F.** 37°C strain BAA-450, **G.** 28°C strain RE-87, **H.** 37°C strain RE-87. Mean and standard deviation of three biological replicates are shown.

### *V. coralliilyticus* biosynthesizes glycine betaine (GB) and uptakes numerous compatible solutes for osmotic protection

Bioinformatics analysis of *V. coralliilyticus* genomes identified the *betIBA* cluster encoding homologs of enzymes required for the biosynthesis of GB from choline and six putative compatible solute transporters in all 35 strains present in the NCBI genome database (as of March 2024). To confirm biosynthesis of GB from choline, ^1^H-nuclear magnetic resonance analysis (^1^H-NMR) was performed with *V. coralliilyticus* cells grown in M9G 4% NaCl with 1 mM choline and from cells grown in the absence of choline. ^1^H-NMR spectral analysis of ethanol extracts did not show any peaks in the absence of choline, but GB peaks were present in CN52H-1 cells (**Fig. 3A-B**) grown in M9G 4% NaCl supplemented with choline and in strains OCN014 (**Fig. S2)** and BAA-450 (data not shown). To determine the compatible solutes that are transported into the cell for osmotic stress protection, the growth of *V. coralliilyticus* strain CN52H-1 was examined in M9G 5% NaCl supplemented with one of ten different compatible solutes (**Fig. 3C**, **Fig. S3**). In the absence of compatible solutes in the media, *V. coralliilyticus* does not grow in M9G 5% NaCl as demonstrated (**Fig. 2**, **Fig. S3)**. The growth pattern of *V. coralliilyticus* cells grown in each compatible solute was monitored for 24 h at 28°C and area under the curve (AUC) was calculated (**Fig. 3C**). Seven of the ten compatible solutes tested allowed growth in M9G 5% NaCl, suggesting they were taken up and used for osmotic stress protection (**Fig. 3C**). The presence of trimethylamine-N-oxide, *myo*-inositol (MI), or glycerol did not allow growth in M9G 5% NaCl (**Fig. S3)**. GB, dimethylsulfoniopropionate (DMSP), ectoine, choline, or dimethylglycine (DMG) rescued growth to the greatest extent reaching a final OD595 ranging from 0.45 to 0.69 (**Fig. S3)**. The compatible solutes γ-amino-butyric acid (GABA) and creatine had longer lag phase times with lower final OD595 of 0.23 and 0.31, respectively indicating that they were less effective as osmolytes (**Fig. S3)**.

**Fig. 3.**
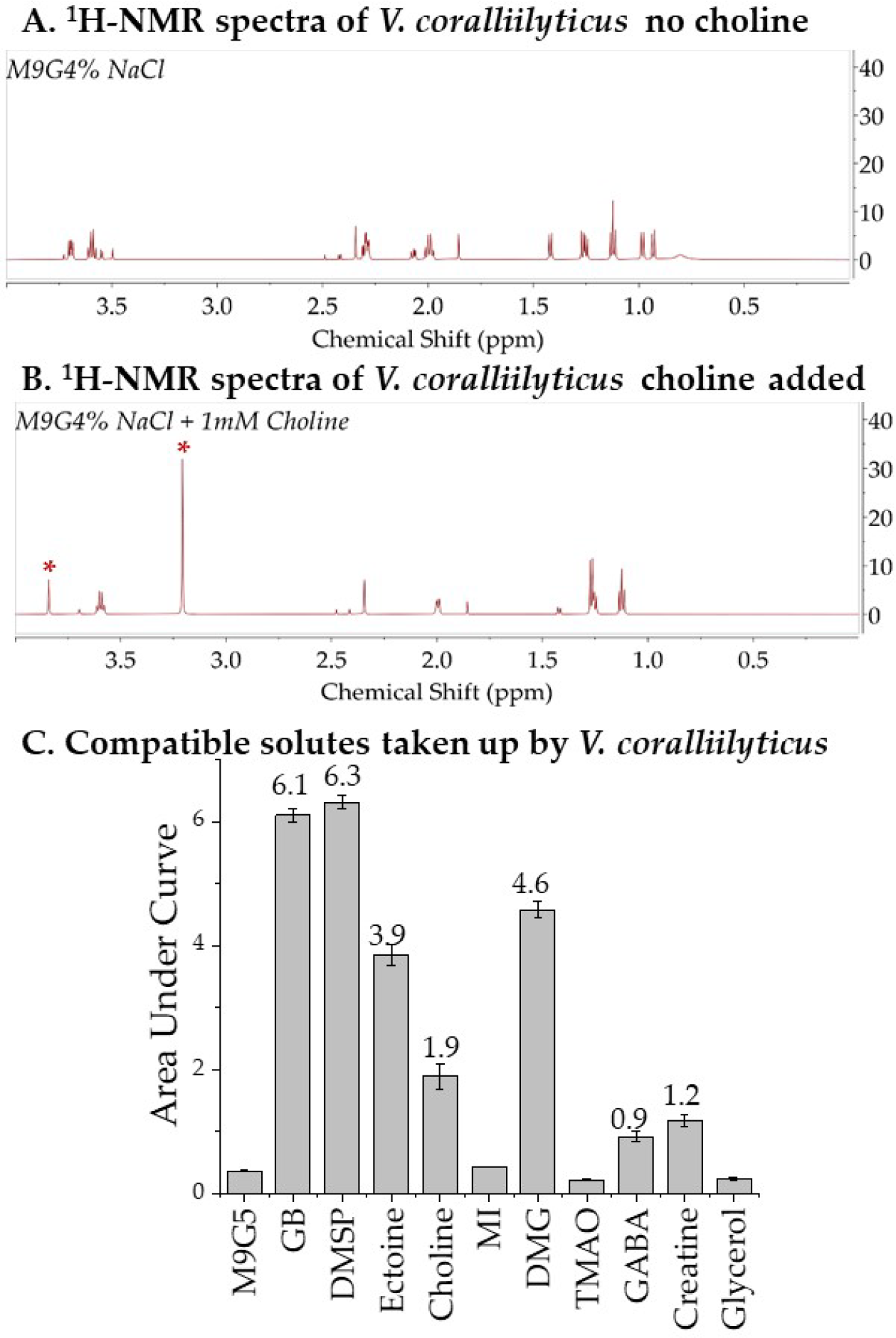
*V. coralliilyticus* compatible solute biosynthesis and uptake systems. **A.** ^1^H-NMR spectra of *V. coralliilyticus* CN52H-1 cellular extract grown in minimal media (M9G) 4% NaCl without choline and **B.** with the addition of 1 mM choline. The spectral peaks corresponding to glycine betaine are labeled with red asterisks. **C.** *V. coralliilyticus* was inoculated into M9G 5% NaCl with 1 mM GB (glycine betaine), DMSP (dimethylsulfoniopropionate), ectoine, choline, MI (*myo*-inositol), DMG (dimethylglycine), trimethylamine N-oxide (TMAO), gamma-aminobutyric acid (GABA), creatine, or glycerol. Growth was measured for 24 h at 28°C and the calculated area under the curve AUC calculated shown above each compatible solute that rescued growth. Mean and standard deviation of three biological replicates are shown.

### *Myo* -inositol catabolism clusters are present in all strains of *V. coralliilyticus* and *V. mediterranei*

Bioinformatics analysis identified putative *myo*-inositol catabolic gene clusters that were present in all but one of the 35 *V. coralliilyticus* and 18 *V. mediterranei,* a species also linked to coral disease, annotated genomes in the NCBI genome database (as of March 2024). However, *V. coralliilyticus* SCSIO 43001 (82) was the sole exception, the average nucleotide identify (ANI) of this strain with BAA-450 is only 87.85 which suggests a misclassification of this strain. Therefore, strain SCSIO 43001 was not included in this study. In *V. coralliilyticus* two distinct gene clusters for the regulation, transport, and catabolism of *myo*-inositol were present (**Fig. 4A**), which we named region 1 and 2. Gene annotation and naming were established based upon amino acid similarity to the *myo*-inositol genes present in *Salmonella enterica*. No synteny was found between *myo*-inositol gene clusters in *V. coralliilyticus* and *S. enterica* or *C. vibrioides* (**Fig. S4)**. Functionally similar genes were present in both region 1 and 2, but shared low sequence similarity (27% to 50%) and lacked gene synteny between each other.

**Fig. 4.**
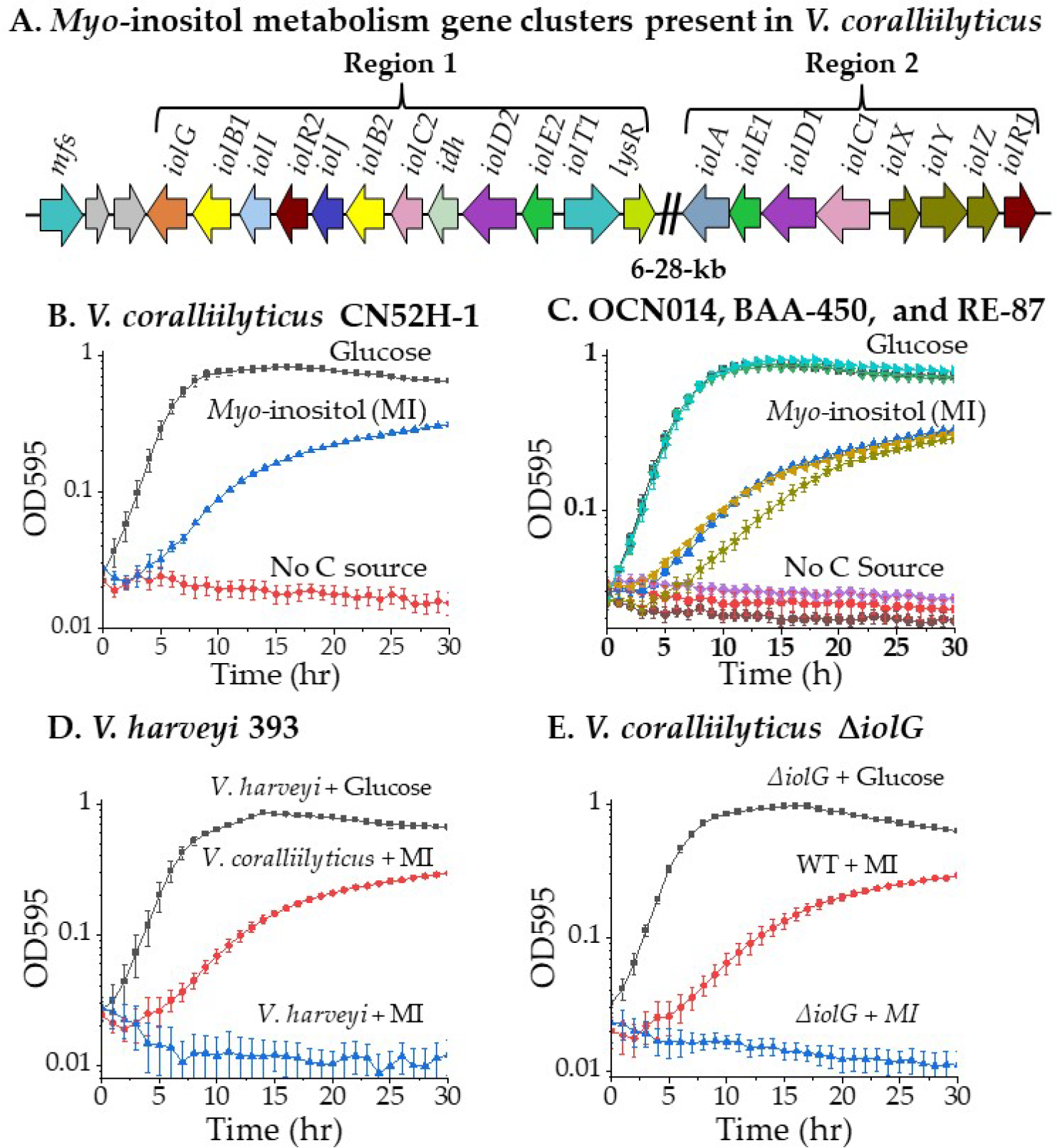
*V. coralliilyticus* utilized *myo*-inositol as a sole carbon source. **A.** Schematic of the gene order of *myo*-inositol transporter, catabolism and regulatory genes in *V. coralliilyticus*. **B.** Growth curve analysis in minimal media (M9 2% NaCl) containing either 20 mM glucose (squares) or 20 mM *myo*-inositol (MI, triangles) or no carbon source (circles) for *V. coralliilyticus* CN52H-1. **C.** *V. coralliilyticus* strains OCN014, BAA-450, and RE-87. **D.** *V. harveyi* 393 and **E.** *V. coralliilyticus* Δ*iolG.* Growth was measured every hour for 30 h at 28°C. Mean and standard deviation of four biological replicates are shown.

Region 1 contained fifteen genes, which included a predicted operon composed of ten genes *iolEDidhiolCBJRIBG* encoding putative enzymes responsible for *myo*- inositol conversion to dihydroxyacetone and malonate semialdehyde (**Fig. 4A** and **Fig. 1**). Gene designations were based on sequence similarity to the *myo*-inositol cluster in *S. enterica* (**Table S1**). Two copies of IolE and IolD are present, a copy in each region. The *iolE* and *iolD* genes encode enzymes involved in the second and third steps of the catabolic pathway. Region 1 copies are named IolE2 and IolD2 and shared 38% amino acid identity to the enzymes from *S. enterica* (**Table S1**). The *iolC* gene encodes a putative 5-dehydro-2-deoxyglucokinase and shared 33% amino acid identify with IolC from *S. enterica*. The next step in the pathway is completed by an oxidoreductase encoded by *iolB.* Two copies of this gene were found within region 1 (named *iolB*1 and *iolB* 2) that shared 63.5% and 31% amino acid identify respectively to IolB from *S. enterica* (**Table S1**). The *iolG* gene encoded an inositol 2-dehydrogenase required for the first step in the pathway. IolG shared 36.1% amino acid identity to IolG2 from *S. enterica*. An *idh* gene within the operon encoded a Gfo/Idh/MocA-type oxidoreductase, which could have a dehydrogenase function to convert isomers of inositol into the degradation pathway. Region 1 also contained two MFS transporters, a LysR family transcriptional regulator, a glycosyltransferase and a polysaccharide biosynthesis protein.

Region 2 contained a predicted operon *iolCDEA,* which shared greater amino acid identity with Iol in *S. enterica* than proteins in region 1, with similarity ranging from 43.7% to 67.4% (**Table S1**). The *iolA* gene encoded a CoA-acylating methylmalonate-semialdehyde dehydrogenase involved in the last step in the pathway (**Fig. 4A**). Divergently transcribed from *iolCDEA* was *iolXYZR* that encoded an ABC-type transporter IolXYZ and an RpiR-type regulator, which we named IolR1 as it shared 67% identity with IolR from *S. enterica* (**Fig. 4A**). *S. enterica* does not possess an ABC-type transporter that uptakes *myo*-inositol, but the substrate binding protein IolX from *V. coralliilyticus* shared 31% amino acid identity with IbpA a known transporter of *myo*- inositol in *C. vibrioides* (79).

A similar gene arrangement in region 1 and 2 was present in the genomes *V. coralliilyticus* strains CN52H-1, OCN014, BAA-450, and RE87 (**Fig. S5**). In *V. mediterranei* (formerly *V. shilonii)*, strains of which are also coral pathogens, a single *iol* cluster is present comprised of 18 genes, putative operons *iolCA, iolXYZR, iolEDidhiolCBJRIBG* and *iolT1lysR* (**Fig. S4).** These data suggest that these strains have the potential to catabolize *myo*-inositol as a sole carbon source. It is of interest to note, that species belonging to the Harveyi clade, specifically 65 *V. harveyi,* 13 *V. campbellii,* 23 *V. owensii,* and 10 *V. jasicida* strains in the NCBI genome database (as of March 2024), contained *myo*-inositol region 1, but not region 2 (**Fig. S6**). That is, no *iolA* or *iolXYZ* homologs are present in these species.

### *Vibrio coralliilyticus* can utilize *myo* -inositol as a sole carbon source

To demonstrate *myo*-inositol catabolism by *V. coralliilyticus* CN52H-1, OCN014, BAA-450, and RE87, strains were grown in M9 2% NaCl media supplemented with 20 mM glucose or 20 mM *myo*-inositol or no carbon source. In M9 2% NaCl supplemented with *myo*-inositol, *V. coralliilyticus* CN52H-1 reached a final OD595 of 0.31 with a lag time of 3 h (**Fig. 4B**). In M9 2% NaCl glucose, there was a 1 h lag phase and a final OD595 of 0.65 was recorded and in the absence of a carbon source, no growth occurred (**Fig. 4B**). Next, we examined the ability of strains OCN014, BAA-450, and RE87 to grow on *myo*-inositol as a sole carbon source. As expected, these strains followed a similar pattern of growth as CN52H-1 on *myo*-inositol or glucose as sole carbon sources (**Fig. 4C**). We then examined a *V. harveyi* strain since it only contained region 1 to determine whether growth on *myo*- inositol can still occur. *V. harveyi* grew in M9 2% NaCl glucose but did not grow in *myo*- inositol as a sole carbon source, whereas the control *V. coralliilyticus* did (**Fig. 4D**). This suggested that region 2 is essential for *myo*-inositol catabolism, likely requiring the *iolA* gene and/or *iolXYZ.* To demonstrate the requirement for region 1 for *myo*-inositol catabolism, an in-frame non-polar deletion of *V. coralliilyticus iolG* (HRJ43_RS06535) was constructed. The *iolG* gene encodes an inositol dehydrogenase essential in the first step of the *myo*-inositol pathway and the last gene in region 1 of the *iol* operon (**Fig. 1**). To demonstrate there was no overall growth defect in the construction of the Δ*iolG* mutant, the strain was grown in M9 2% NaCl glucose, which showed a growth pattern identical to the wild type strain (**Fig. 4E**). As expected, the Δ*iolG* strain showed no growth in M9 2% NaCl *myo*-inositol as a sole carbon source, with the wild type strain used as a positive control (**Fig. 4E**). Overall, the data showed that *V. coralliilyticus* has a functioning *myo*-inositol pathway and both regions 1 and 2 are required for catabolism.

### Ligand modeling predicts *V. coralliilyticus* IolT1 and IolXYZ are *myo* -inositol transporters

Two major facilitator superfamily (MFS)-type transporters are present within region 1 and one ABC-type transporter IolXYZ within region 2. Ligand modeling was performed by comparison to known transporters of *myo*-inositol to predict which of these are likely responsible for *myo*-inositol transport. The MFS transporter from *B*.

*subtilis* IolT (WP_003234027.1) and *S. enterica* IolT1 (NP_463279.1) were selected as these transporters were previously shown to transport *myo*-inositol (63). The MFS protein (WP_099607352.1) within the region 1 cluster shared 48.8% amino acid identity to that of *S. enterica* IolT1, which we named IolT1. Whereas the MFS protein (WP_172853823.1) outside of the cluster shared only 25% similarity to that of *S. enterica,* which we named MFS. Docking of *myo*-inositol with *B. subtilis* IolT and *S. enterica* IolT1 predicted a mode of interaction with seven and five putative binding residues, respectively, and favorable binding free energy changes (ΔG) of −7.0 kcal/mol and −6.1 kcal/mol. *Myo*-inositol docking with *V. coralliilyticus* IolT1 predicted a favorable ΔG of −6.2 kcal/mol and four putative binding residues. MFS-type transporters are known to coordinate substrate binding with transmembrane (TM) domains TM7 and TM10 (83). Alignment of *B*.

*subtilis* IolT, *S. enterica* IolT1, *V. coralliilyticus* IolT1 and *V. coralliilyticus* MFS was performed, and TM domains were identified. Predicted binding residues found in the three models are shown with blue (*V. coralliilyticus* IolT1), green (*S. enterica* IolT1), and orange (*B. subtilis* IolT) arrows, which were absent from MFS (**Fig. S7).** The two glutamine (Q273, Q274) and asparagine (N279) in TM7 are found in all three modeled sequences. In TM10 three of the four sequences also contained a conserved tryptophan (W377) residue. All three models predicted a residue that aligns at Y368. Both *V. coralliilyticus* IolT1 and *S. enterica* IolT1 contain a conserved tyrosine (Y) residue here, but *B. subtilis* IolT contained glutamine (Q) (**Fig. S7).** Taken together this similarity in the binding pockets of these transporters suggested that *V. coralliilyticus* IolT1 is likely able to transport *myo*-inositol.

The ABC-type transporter substrate binding protein IbpA from *C. vibrioides* had previously been shown to bind *myo*-inositol (79) and IolX (WP_227495657.1) from *V. coralliilyticus* has 87% coverage and 31% amino acid identity to IbpA. Ligand docking analysis of *myo*-inositol with IbpA and IolX predicted favorable modes of action with ΔG of −7.4 kcal/mol and -5.3 kcal/mol, respectively. Putative binding residues were identified for each substrate binding protein and are presented in the amino acid alignment (**Fig. S8**). The green arrows indicate IbpA predicted binding residues and blue arrows indicate IolX predicted binding residues. Five aligned residues were predicted to be important in both docking models: 63 aspartic acid (D)/ glutamine (Q), 141 aspartic acid (D)/ asparagine (N), 187 glutamine (Q)/ serine (S), 269 aspartic acid (D), and 289 glutamine (Q) (**Fig. S8B**). Based upon the favorable docking model and select conserved residues in the alignment with a known *myo*-inositol transporter, IolXYZ is predicted to be capable of *myo*-inositol transport.

### The MFS transporter IolT1 and ABC-type transporter IolXYZ are responsible for *myo* -inositol uptake for catabolism

To evaluate the ability of IolT1 and IolXYZ to transport *myo*-inositol, in-frame non-polar deletion mutants were generated in the *iolT1* (HRJ43_RS06585) and *iolX (HRJ43_RS06670)* genes and a double deletion mutant *iolT1/iolX* was also constructed. To show that these mutants contained no overall growth defects, all three mutants and wild type were each grown in M9 2% NaCl supplemented with 20 mM glucose. All three mutants showed an identical growth pattern to wild type indicating that the mutants do not contain any additional defects (**Fig. 5A**).

**Fig. 5.**
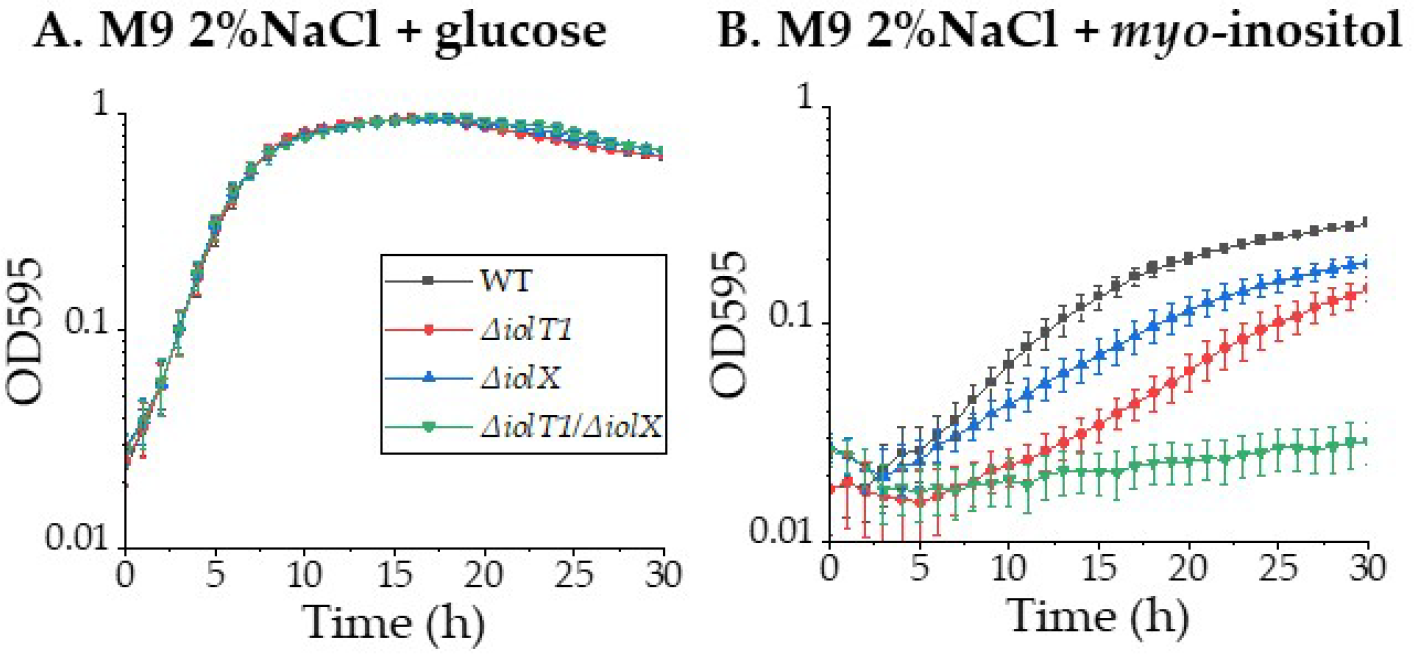
Growth pattern analysis of *V. coralliilyticus myo*-inositol transporters deletion mutants. Growth analysis of wild type, Δ*iolT1*, Δ*iolX*, and Δ*iolT1*/ Δ*iolX* in minimal media (M9 2% NaCl) with **A.** 20 mM glucose or **B.** 20 mM *myo*-inositol as a sole carbon source. Growth was measured every hour for 30 h at 28°C. Mean and standard deviation of three biological replicates are shown.

Next, the three mutants and wild type were each grown in M9 2% NaCl supplemented with 20 mM *myo*-inositol for 30 h at 28°C (**Fig. 5B**). Under these growth conditions, the Δ*iolT1* mutant had a decreased maximum growth rate of 0.08 h^-1^ compared to wild type; 0.19 h^-1^ and reached a lower final OD595 (0.15) as compared to wild type (0.29) (**Fig. 5B**). The Δ*iolT1* mutant also showed a lag time of 6 h compared to 3 h for wild type (**Fig. 5B** and **Fig. S9**). These data indicated that IolT1 is utilized for the transport of *myo*-inositol, but another transporter also transports this compound into the cell. Growth pattern analysis of the Δ*iolX* mutant also showed a decreased growth rate of 0.11 h^-1^ and a lower final OD595 of 0.19 compared to wild type, but no difference in the lag phases (**Fig. 5B** and **Fig. S9**). These data show that IolXYZ is also a *myo*- inositol transporter. Next, we examined the double deletion mutant *iolT1/iolX* to determine whether any other transporters could uptake *myo*-inositol. The double mutant failed to grow in M9 supplemented with *myo*-inositol as a sole carbon source showing *V. coralliilyticus* has two *myo*-inositol transporters (**Fig. 5B**). The differences between the lag time of the single mutants, 3 h lag when only *iolT1* was present and a 6 h lag for when only *iolX,* indicated that IolT1 is likely more efficient at *myo*-inositol uptake compared to IolXYZ (**Fig. 5B** and **Fig. S9**).

### Phylogenetic analysis of IolG from region 1 and IolA from region 2 among *Vibrionaceae* reveals different evolutionary histories

The low sequence similarity between comparable proteins that are present in both region 1 and 2, and the lack of synteny between the regions suggest different evolutionary histories. To examine this further, a phylogeny based on the IolG protein from region 1 and IolA from region 2 were constructed among members of the family *Vibrionaceae*(**Fig. 6** and **Fig. S10A**). In addition, a phylogeny based on the housekeeping protein RpoB was also constructed among the same set of species to represent their ancestral relationships and the major clades to which each species belongs (**Fig. S10B).** To accomplish this, BLAST analysis was first performed using IolG (WP_038162996.1) as a seed to identify homologous protein sequences in the NCBI genome database. A total of 53 sequences with >95% query coverage and >60% amino acid similarity were downloaded and aligned using CLUSTALW (84). The genome context of these IolG proteins within each species was determined to confirm the presence of additional *myo*-inositol metabolism genes. Phylogenetic trees were constructed, and a representative IolG tree is shown in Figure 6A constructed using the neighbor-joining method based on evolutionary distance calculated using the JTT matrix-based method (85). IolG was present in 32 *Vibrio,* 11 *Photobacterium*, 6 *Enterovibrio* and 4 *Grimontia* species (**Fig. 6A**). Among *Vibrio* species, the IolG proteins were found dispersed within three major clusters on the tree. The *V. coralliilyticus* IolG, a species that belongs in the Coralliilyticus clade, branched with high bootstrap value with members of the Mediterranei clade (**Fig. 6A**). Branching distantly from these species was a highly similar cluster of IolG proteins indicated by short branch lengths in species from the Harveyi clade (**Fig. 6A**). Within this tight cluster was IolG from *V. vulnificus,* a species that is not a member of the Harveyi clade (**Fig. 6A** and **Fig. S10B**). These data indicated that IolG and by extension region 1 in *V. vulnificus* was acquired by horizontal gene transfer from a Harveyi clade species. Species in the Harveyi clade did not contain region 2 or a homolog of IolA. The next large clustering of IolG from *Vibrio* species was from four species of the Splendidus clade and branching distantly from these were IolG from nine *Photobacterium* species. Branched distantly from this group were IolG from two *Photobacterium* species that clustered divergently from species from the Cholerae and Splendidus clades (**Fig. 6A** and **Fig. S10B**). Along with the IolG phylogeny, the differences in gene order and gene content among the *Vibrio* species suggest the *iol* gene clusters have different evolutionary histories and were likely acquired multiple times within the genus (**Fig. 6B**). The IolG proteins from five *Grimontia* species clustered closely together with short branch lengths and identical cluster indicating a single acquisition within the genus (**Fig. 6B**). IolG from species of the genus *Enterovibrio* branched divergently from one another (**Fig. 6A**). Comparisons between the IolG and RpoB trees show that the IolG tree does not follow species evolutionary histories with the exception of *Grimontia* and *Enterovibrio* and indicate that IolG was acquired by horizontal transfer in *Vibrio* and *Photobacterium* species **(Fig. S10B**).

**Fig. 6.**
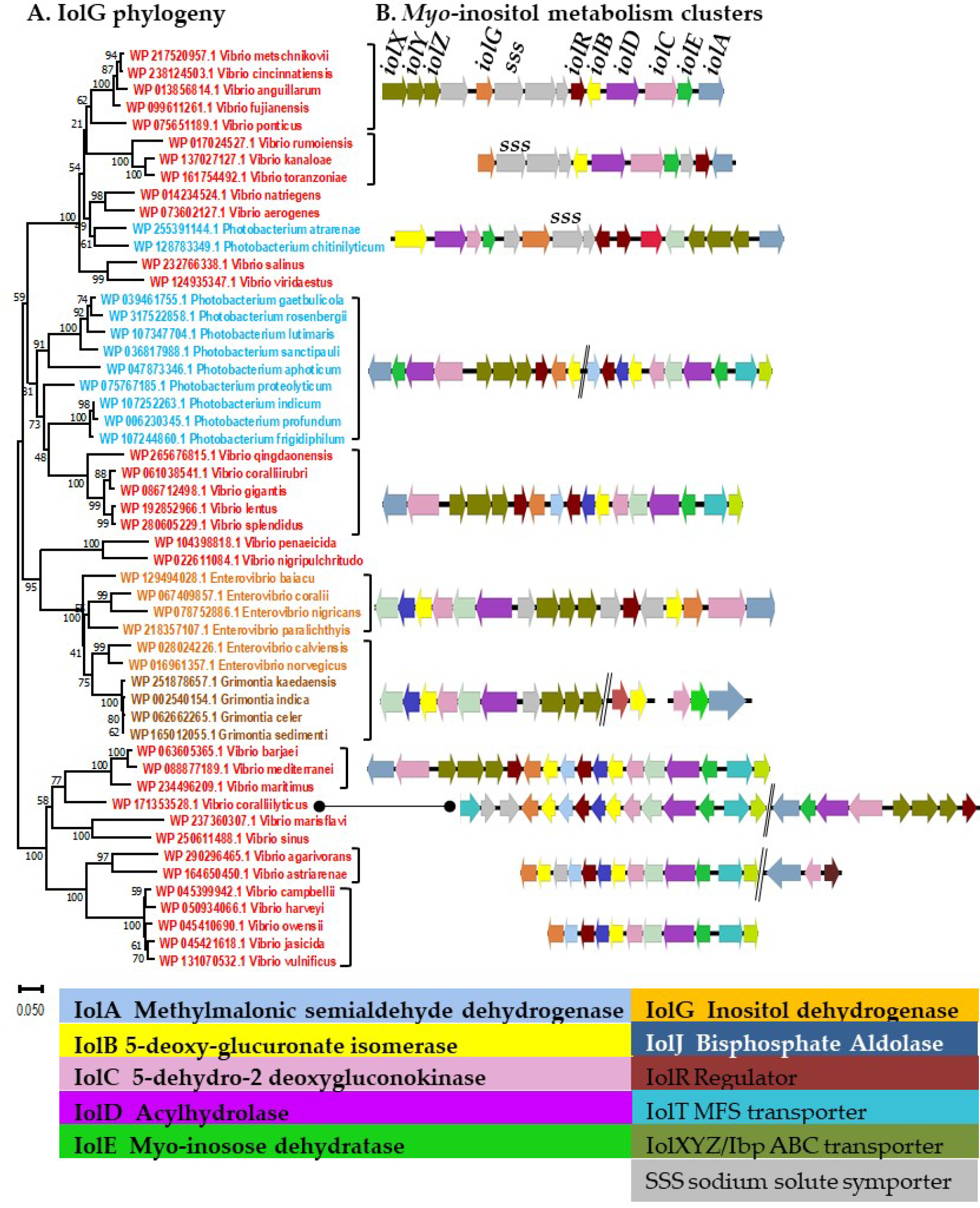
Phylogenetic distribution of IolG and Iol cluster organization among *Vibrionaceae*. **A.** IolG evolutionary history was inferred using the Neighbor-Joining method (118) with bootstrap value of 1000 (123). The optimal tree with the sum of branch length = 4.11930205 is shown. The tree is drawn to scale, and evolutionary distances were computed using the JTT matrix-based method (85) with units of the number of amino acid substitutions per site. Shown are 53 amino acid sequences, ambiguous positions were removed for each sequence pair (pairwise deletion option) with a total of 332 positions. Evolutionary analyses were conducted in MEGA X (117). IolG was present in only 32 *Vibrio* (red), 11 *Photobacterium* (blue), 6 *Enterovibrio* (orange), and 4 *Grimontia* (brown) species. **B**. Iol cluster organization among *Vibrionaceae* species. Arrows represent open reading frame and direction of transcription, double black lines represent genes between regions.

The IolA protein, a CoA-acylating methylmalonate-semialdehyde dehydrogenase, was selected as the representative of region 2 and used as a seed for BLAST analysis to identify homologs among the *Vibrionaceae*. This analysis uncovered eight *Vibrio* species that contained IolG that did not have a copy of IolA or any of the proteins present in region 2 (**Fig. S10A**). These included species from the Harveyi clade mentioned above as well as *V. astriarenae, V. agarivorans, V. ponticus*, and *V. vulnificus.* In the IolA tree, the bootstrap values for internal branch nodes for most *Vibrio* and *Photobacterium* species were low and indicated that the relationships of IolA from these species is not clear (**Fig. S10A**). In contrast, the bootstrap values for IolA from *Grimontia* species were robust with short branch lengths and indicated that they are highly related to one another (**Fig. S10A**). Branched divergently from these were IolA from *Enterovibrio* species with greater than 85% confidence in many external nodes and IolA from *V. penaeicida* and *V. nigripulchritudo* also clustered distantly within this group similar to the IolG tree (**Fig. S10A**). The position of IolA from *Photobacterium* species within the tree could not be determined as the internal bootstrap values were < 40%. The IolA protein from *Photobacterium chitinilyticum* branched distantly with IolA from *Vibrio* species similar to IolG from this species. In most *Vibrio* species, divergent branches with long-branch lengths were present, which represented low IolA similarity to each other. Comparisons between the IolG and IolA trees and comparative *iol* cluster analysis suggest that region 1 and 2 were acquired separately, each with a different origin and ancestral history (**Fig. 6** and **Fig. S10A**). The phylogeny and *iol* cluster comparisons demonstrate a polyphyletic origin for the *myo*-inositol metabolic clusters likely acquired via horizontal transfer. However, there were no signatures of recent horizontal transfer such as the presence of IS elements, transposases, integrases, phage genes, or plasmid genes within 10 genes of the metabolic gene clusters (with the exception of *V. rumoiensis,* which contained an integrase adjacent to *iolA)*.

To examine *myo*-inositol metabolism evolutionary history further, an IolG tree was constructed from IolG homologs outside of the *Vibrionaceae* family (**Fig. 7A**). Within this tree, IolG from *Vibrio* species clustered most closely with *Plesiomonas shigelloides*and *Aeromonas* species, both pathogens of fish and humans and clustering divergently from these was IolG from *Providencia* species. A common denominator in all these groups was an 11-13 gene cluster that contained a sodium solute symporter (SSS) type transporter. The *iol* cluster was present in a limited number of strains of *A. allosaccharide*, *A. dhakensis*, *A. finlandensis, A. hydrophila, A. media, A. rivopollensis, A. simiae,* and *A. taiwanensis.* In *A. dhakensis* strains, also an important pathogen of fish, the *iol* genes were present at a tRNA-Ser locus, within a region that contained phage genes and genes for a restriction modification system (**Fig. 7B**). In *A. media, A. allosaccharophila,* and *A. finlandersis,* the *iol* gene cluster was also present adjacent to the same tRNA-Ser locus, but had different flanking genes and was present in one strain of each species.

**Fig. 7.**
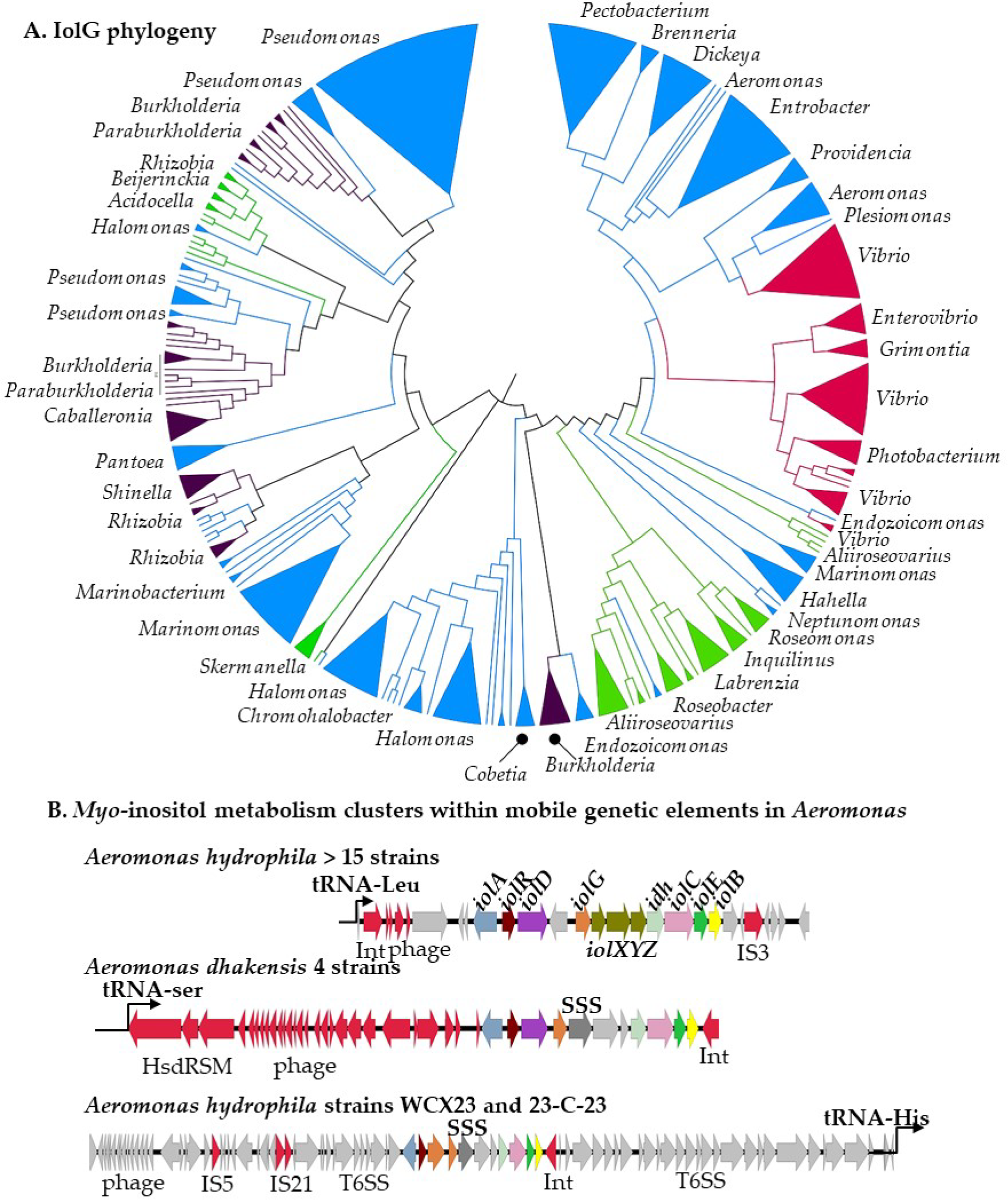
IolG phylogeny among marine species. Condensed consensus tree with collapsed branches is shown, original tree is shown in **Fig. S11**. IolG phylogeny was inferred using the Neighbor-Joining method (118). This analysis involved 358 amino acid sequences. Blue and red collapsed branches represent Gamma-, purple represents Beta-, and green represents Alpha-Proteobacteria. **B.** Genome context of *myo*-inositol metabolism clusters within mobile genetic elements in *Aeromonas* species.

In a separate group of *A. hydrophila* strains (>15 strains in the NCBI genome database), the *iol* gene cluster contained the *iolXYZ* transporter. The IolG from these *A. hydrophila* strains clustered divergently from other *Aeromonas* species and branched distantly with *Enterobacter* species. These *A. hydrophila* strains were previously characterized as hypervirulent fish pathogens. The *iol* cluster in these strains was inserted adjacent to a tRNA-Leu locus and was surrounded by integrase, transposase, and phage genes, markers of horizontal gene transfer (**Fig. 7B**). Branching divergently from these were IolG from species of the genera *Pectobacterium (formerly Erwinia species), Brenneria,* and *Dickeya* (**Fig. 7A**). Genomic analysis of *Brenneria*, *Dickeya,* and *Pectobacterium*showed that the *myo*-inositol metabolic genes were present in a single conserved cluster; in *Pectobacterium* and *Brenneria* the operon structure was *iolADGXYZ_idh_iolCEB* with an ABC-type transporter (*iolXYZ)*. In *Dickeya* species, the region was similar to the region in *Pectobacterium* with the exception of three genes inserted between *iolD* and *iolG.* In *Pectobacterium* species, the *iol* cluster was flanked by an addiction system (toxin-antitoxin genes) and the region was located near a tRNA-Asn locus (**Table S2-S5**). In *Brenneria izbisi,* the *iol* region was also located at the same tRNA-Asn locus, but in this species the cluster was flanked by a T6SS and an integrase (**Table S6**). For *Dickeya* and *Pectobacterium*, all species in the IolG tree showed short branch lengths indicating these proteins are highly related to one another within each genus suggesting IolG is ancestral to the genera. This indicates that the region was acquired once, which is also evidenced by their presence at the same genome location (tRNA-Asn).

The discovery of inositol catabolism among a relatively small group of *Vibrio* species made us wonder about the distribution of these genes among marine bacteria in general. Previous work by Fuchs and colleagues indicated that *iol* gene clusters were present in over 30 bacterial genera associated with aquatic environments (60). To investigate this further, we examined bacteria associated with different marine hosts for the presence of IolG. This analysis identified *iol* clusters in *Aliiroseovarius, Burkholderia, Caballeronia, Cobetia, Endozoicomonas, Hahella, Halomonas, Marinomonas, Marinobacterium, Labrenzia, Marinomonas, Marinobacter, Oceanspirallaceae,Paraburkholderia Pseudomonas,* and *Rhizobia* amongst others (**Fig. 7A**). The IolG phylogeny of these groups showed divergent sequences among closely related species, with species from Alpha-, Beta- and Gamma-Proteobacteria clustered together (**Fig. 7A**). Some genera showed tight clustering of IolG with short branch lengths, such as *Hahella, Halomonas, Marinomonas,* and *Pseudomonas* (**Fig. S11**). Whereas *Burkholderia (*previously *Pseudomonas), Caballeronia,* and *Paraburkholderia* species, all members of the *Burkholderiaceae* family, clustered with each other with short branch lengths and in some cases grouped with *Pseudomonas* species (**Fig. S11**). These genera as well as being significant human and animal pathogens are also characterized as symbionts of protozoa, cnidarians, insects, and plants (86–89). Within the groups that contained an *iol* cluster in our analysis were many genera isolated from reef sponges, corals, marine algae, crustaceans, molluscs and marine fish. For example, *Endozoicomonas, Halomonas, Inquilinus, Labrenzia, Roseobacter, Pseudomonas, Photobacterium,*and *Vibrio* are known symbionts or present in the holobiont of corals (**Fig. S11**) (10, 90, 91). Additionally comparative analysis of the cluster showed common themes such as the presence of a single cluster within which was invariably the ABC-type transporter IolXYZ. A common variation among these clusters was the presence or absence of IolA within the cluster or its presence elsewhere on the genome (**Fig. S12**). Additionally, a difference between these *iol* clusters in marine species compared to the clusters in Enterobacterales (*Brenneria, Dickeya, Enterobacter, Pectobacterium*) was the presence of multiple isomerases and dehydrogenases genes (**Fig. S12**). For example, in *Burkholderia cepacia* four additional dehydrogenase and isomerases were present within the cluster and *Halomonas titanicae*, three additional dehydrogenases and isomerases were present (**Fig. S12**).

## DISCUSSION

Our examination of *V. coralliilyticus* response to changes in osmotic stress under different growth temperatures demonstrated the importance of NaCl for optimal growth and growth at high temperatures. The data showed that *V. coralliilyticus* can grow in 1% - 4% NaCl, but grows optimally in 2% NaCl at 28°C and has an absolute requirement for NaCl for growth at 37°C. Compared to other *Vibrio* species such as *V. parahaemolyticus* that grow in up to 7% NaCl without the addition of compatible solutes, *V. coralliilyticus* has a relatively low salinity tolerance of 4% NaCl (92–95). A major difference between *V. coralliilyticus* and *V. parahaemolyticus* in their osmotic stress response mechanisms is the absence of the ectoine biosynthesis cluster *ectABC_asp,* which is required for *de novo* biosynthesis of ectoine. Deletion of the ectoine genes in *V. parahaemolyticus* inhibited growth in NaCl concentration above 5% NaCl in the absence of exogenous osmolytes (95). *V. coralliilyticus* biosynthesized GB from exogenous choline and transported into the cell a wide range of osmolytes including choline, GB, DMSP, DMG, and ectoine for osmotic protection similar to *V. parahaemolyticus* (92, 93). *Myo*-inositol is a compatible solute more commonly associated with a few hyperthermophile bacteria as well as many eukaryotic species such as algae and plants (96–99). Therefore, it came as no surprise that *V. coralliilyticus* did not use *myo*-inositol for osmotic protection. However, our bioinformatics analysis did identify a genomic region on chromosome 2 in *V. coralliilyticus* that contained *myo*-inositol catabolism gene clusters. This region was present in all *V. coralliilyticus* genomes in the NCBI genome database showing it is a conserved phenotype in this species. Our analysis showed that *V. coralliilyticus* strains can utilize *myo*-inositol as a sole carbon source with a 3 h lag phase. The short lag phase is meaningful because previous studies on *S. enterica* have shown a lag time of over 40 hours (67, 69, 70, 73). *Caulobacter vibrioides* (68, 79), *Rhizobium leguminosarum* (formerly *trifolii)* (74), and *Sinorhizobium meliloti* (100) were also shown to have long lag times of about 14 h when grown on *myo*-inositol as a sole carbon source. A possible reason for this efficient use of *myo*-inositol by *V. coralliilyticus* is the presence of two transporters for uptake into the cell, IolT1 and IolXYZ. As shown in the deletion mutant analysis of these systems, both have somewhat similar *myo*- inositol uptake affinities. Additionally, the regulation of the catabolism genes is likely very different in these divergent species and also control of transport may be different. Our analysis also showed that *V. harveyi* could not utilize *myo*-inositol as a carbon source and this is likely due to the absence of region 2, and IolA specifically. Whether these strains contained region 2 and lost it or never obtained the region is hard to determine but given the high similarity of region 1 among these species the most parsimonious explanation is that region 2 was never acquired. It is possible that the region in *V. harveyi* could be used to metabolize derivatives of inositol.

We speculate that during periods of thermal stress, the increased production of inositols by the coral and/or their endosymbiotic algae, as has been shown by Hillyer and colleagues (53), is a metabolic opportunity for bacteria. The ability to consume readily available inositols could provide a selective advantage for *Vibrio* commensal or opportunistic pathogens to increase in abundance within the coral microbiome. In turn, consumption of *myo*-inositol might result in depleting the availability of *myo*-inositol as a stress protectant for algae, which could then lead to a decline in algal survival and abundance. In addition, *myo*-inositol is used as a building block in various signal transduction pathways, and the ability of *V. coralliilyticus* to catabolize this sugar could pose a means for signaling interference. Utilization of predominantly a eukaryotic metabolite could be a side consequence that gives *Vibrio* species a competitive advantage over members of the coral microbiome or other eukaryotic host microbiomes that allows the bacterium to exploit a novel food source and enhance colonization. This competitive advantage could be enhanced during disruption of marine host-microbiota interactions due to thermal, salinity, and/or acidic stresses that change microbial diversity and abundance, similar to the human gut microbiome and human health.

Examination of the evolutionary history of IolG showed that the *iol* gene cluster was acquired multiple times within *Vibrionaceae* and given the significant divergence among species, these acquisitions were likely evolutionarily ancient events. Comparison of the IolG and IolA trees showed they were not congruent indicating the two regions were acquired separately. The closest relative to IolG outside of the Vibrionales were a few members of the Aeromonadales (*Aeromonas*) and many Enterobacterales from the genera *Brenneria*, *Dickeya, Enterobacteria*, *Pectobacterium, Plesiomonas, Providencia*. The history of the region among these groups was very different, with signatures of horizontal transfer present among *Aeromonas* species. *Aeromonas* are found in fresh and marine water environments and include many species pathogenic to fish and humans. Two different *myo*-inositol clusters were present in *Aeromonas* species, each containing a single cluster. IolG from seven *Aeromonas* species and from *Plesiomonas shigelloides* branched with IolG from *Vibrio* species and the *iol* cluster in these species encoded a sodium solute symporter (SSS). *P. shigelloides* is the sole member of the genus and is predominantly isolated from aquatic environments and their inhabitants. *P. shigelloides* is presently within the family *Enterobacteriaceae,*but was previously a member of the family *Vibrionaceae*and then *Aeromonadaceae*and has been described as a fish and human pathogen (101). The *iol* cluster is present in all *P. shigelloides* genomes in the NCBI database (as of March 2024) and this species has been shown to utilize *myo*- inositol as a carbon source (101). Included in this IolG group were two *A. hydrophila* strains WCX23 and 23-C-23, where the *iol* metabolic cluster was flanked on both ends by genes encoding a type six secretion system (T6SS) and was inserted next to a tRNA-His locus (**Fig. 7B**). In *A. dhakensis* and the other five species, the *iol* cluster was located adjacent to a tRNA-Ser locus and flanked by signatures of mobile elements (**Fig. 7B**). In hypervirulent *A. hydrophila* strains associated with epidemic disease in fish aquaculture farms in China and the USA, the single *iol* cluster contained the IolXYZ ABC-type transporter and clustered with IolG from *Enterobacter* species. In these hypervirulent *A. hydrophila,* the *iol* cluster was flanked by an integrase and phage genes on one end and an IS3 element on the opposite end and was inserted at a tRNA-Leu locus. Previous studies showed that hypervirulent *A. hydrophila* strains all contained an *iol* cluster, could utilize *myo*-inositol as a sole carbon source, and was an important trait differentiating these strains from other *A. hydrophila* isolates (102–104). *Enterobacter* species are ubiquitous in nature, present in soil and water, and are well known plant pathogens and opportunistic pathogens of humans (105). Several species of *Enterobacter* have been shown to utilize *myo*-inositol as a carbon source (105). Nested within IolG from *Enterobacter* species was IolG from a single *A. taiwanensis* strain. However, the genome for this species was incomplete so the genome context could not be explored further. Overall, the data suggest that in *Aeromonas,* the *iol* gene cluster is a recent addition acquired multiple times on mobile genetic elements and may have contributed to the emergence of hypervirulent *Aeromonas* fish pathogens. Distantly related to these was IolG from *Pectobacterium, Brenneria,* and *Dickeya* species which are broad host range pathogens of plants with a worldwide distribution (**Fig. 7**) (106, 107). The *iol* cluster was universally present in these genera suggesting it is a key phenotype of the group. Emerging evidence from plant rhizosphere studies recently demonstrated that plant-produced inositol catabolized by bacteria correlated with their higher root colonization (108, 109). Whether the presence of *myo*-inositol catabolism genes aids in pathogen’s colonization of plants or is a beneficial trait picked up to take advantage of plant disease releasing inositols remains to be determined. Our analysis also showed that the *iol* cluster is present among a diverse range of marine species (**Fig. S11**). This is in agreement with Fuchs and colleagues demonstrating that over 30 bacterial genera from aquatic environments contained *iol* clusters (60). Many of these species are associated with marine hosts including endosymbionts of corals (110, 111). The distribution of the *iol* cluster within many genera was not universal, suggesting a possible role in niche partitioning.

## MATERIALS AND METHODS

### Bacterial strains, media, and culture conditions

All strains and plasmids used in this study are listed in **Table 1**. Four *Vibrio coralliilyticus* strains were examined in this study, strains CN52H-1, OCN014, BAA-450, and RE87. Unless stated otherwise, *V. coralliilyticus* was grown at 28°C in either lysogeny broth (LB) (Fisher Scientific, Fair Lawn, NJ) with 2% (wt/vol) NaCl (LB 2%NaCl) or M9 minimal media (47.8 mM Na2HPO4, 22 mM KH2PO4, 18.7 mM NH4Cl, 8.6 mM NaCl; Sigma Aldrich) supplemented with 2 mM MgSO4, 0.1 mM CaCl2, 2% (wt/vol) NaCl and 20 mM glucose as the sole carbon source (M9G 2% NaCl). *Escherichia coli* strains were grown in LB supplemented with 1% (wt/vol) NaCl (LB1%) at 37°C. The strain *E. coli* β2155 *λpir* is a diaminopimelic acid (DAP) auxotroph and was grown supplemented with 0.3 mM DAP (112). The *V. harveyi* strain used for analysis was strain 393 (113). Chloramphenicol (Cm) was added to media at a concentration of 12.5 µg/mL when specified.

**Table 1.** Strains and plasmids used in this study.

| Strain/plasmid | Genotype or description | Reference/source |
| --- | --- | --- |
| *Vibrio coralliilyticus* |  |  |
| CN52H-1 | Environmental isolate from reef coral | (16) |
| OCN014 | Pathogen of the coral *Acropora cytherea* | (23) |
| BAA-450 | Pathogen of the coral *Pocillopora damicornis* | (19) |
| RE87 (00-90-6) | Isolated from diseased Pacific oyster larvae | (80, 81) |
| $\Delta iolG$ | Deletion of HRJ43_RS06535 from CN52H-1 | This study |
| $\Delta iolT1$ | Deletion of HRJ43_RS06585 from CN52H-1 | This study |
| $\Delta iolX$ | Deletion of HRJ43_RS06670 from CN52H-1 | This study |
| $\Delta iolT1/\Delta iolX$ | *iolX* deletion in $\Delta iolT2$ mutant background | This study |
| *Vibrio harveyi* |  |  |
| 393 | Isolated from barramundi in Australia | (113) |
| *Escherichia coli* |  |  |
| DH5α λpir | | ThermoFisher Scientific |
| B2155 λpir | $\Delta dapA::erm$ pir for bacterial conjugation | (112) |
| **Plasmids** |  |  |
| pDS132 | Suicide plasmid; CmR; *sacB*, R6Kg ori | (116) |
| pDS*iolG* | pDS132 harboring truncated *iolG* allele | This study |
| pDS*iolT1* | pDS132 harboring truncated *iolT1* allele | This study |
| pDS*iolX* | pDS132 harboring truncated *iolX* allele | This study |

### Osmotic Stress Growth Analysis

*Vibrio coralliilyticus* was grown overnight in M9G 1% NaCl at 28°C with aeration. A 2% inoculation of the overnight culture into fresh M9G 1% NaCl was grown to OD595 of ∼ 0.50 at 28°C with aeration. Exponential cultures were inoculated 1:40 into 200 µL fresh M9G media containing 1% to 6% NaCl. When using compatible solutes glycine betaine (GB), dimethylsulfoniopropionate (DMSP), ectoine, choline, dimethylglycine (DMG), trimethylamine N-oxide (TMAO), gamma- aminobutyric acid (GABA), creatine, *myo*-inositol (MI), and glycerol were used at a final concentration of 1 mM in M9G 5% NaCl. Plates were incubated at 28°C with intermittent shaking for 1 min every hour. Optical densities were taken at 595 nm every hour for a total of 24 h using a Tecan Sunrise microplate reader and Magellan plate reader software (Tecan Systems Inc., San Jose, CA).

### Deletion Mutant Construction

In-frame non-polar deletion mutants of *iolG* (HRJ43_RS06535), *iolT1* (HRJ43_RS06585), and *iolX* (HRJ43_RS06670) were constructed using SOE PCR and allelic exchange as previously described (114). Truncations of the genes of interest, *iolG* (18-bp of the 984-bp gene), *iolT1* (30-bp of the 1452-bp gene), and *iolX* (24-bp of the 1008-bp gene) were generated using the primers listed in **Table 2**. The truncated gene products were ligated with the suicide vector pDS132 using the Gibson assembly protocol and then transformed into *E. coli* Dh5α λpir (115, 116). The plasmids pDSΔ*iolG,* pDSΔ*iolT1,* and pDSΔ*iolX* were then purified and transformed into *E. coli* β2155 *λpir* (DAP auxotroph). Then conjugation followed by homologous recombination into the *V. coralliilyticus* CN52H-1 genome was completed. To select for the single crossover of the plasmids into the genome, plating onto chloramphenicol (Cm) was used and colonies were screened via PCR for the truncated allele. To induce a double- crossover event, the single-cross strains were grown overnight in the absence of Cm. The cultures were then spread on 10% sucrose plates, and healthy colonies were screened for gene truncation, and plasmid excision. Colonies that still contained the plasmid would appear soupy on the 10% sucrose plates because of the presence of *sacB* selectable marker. In-frame deletions were confirmed with sequencing.

**Table 2.**
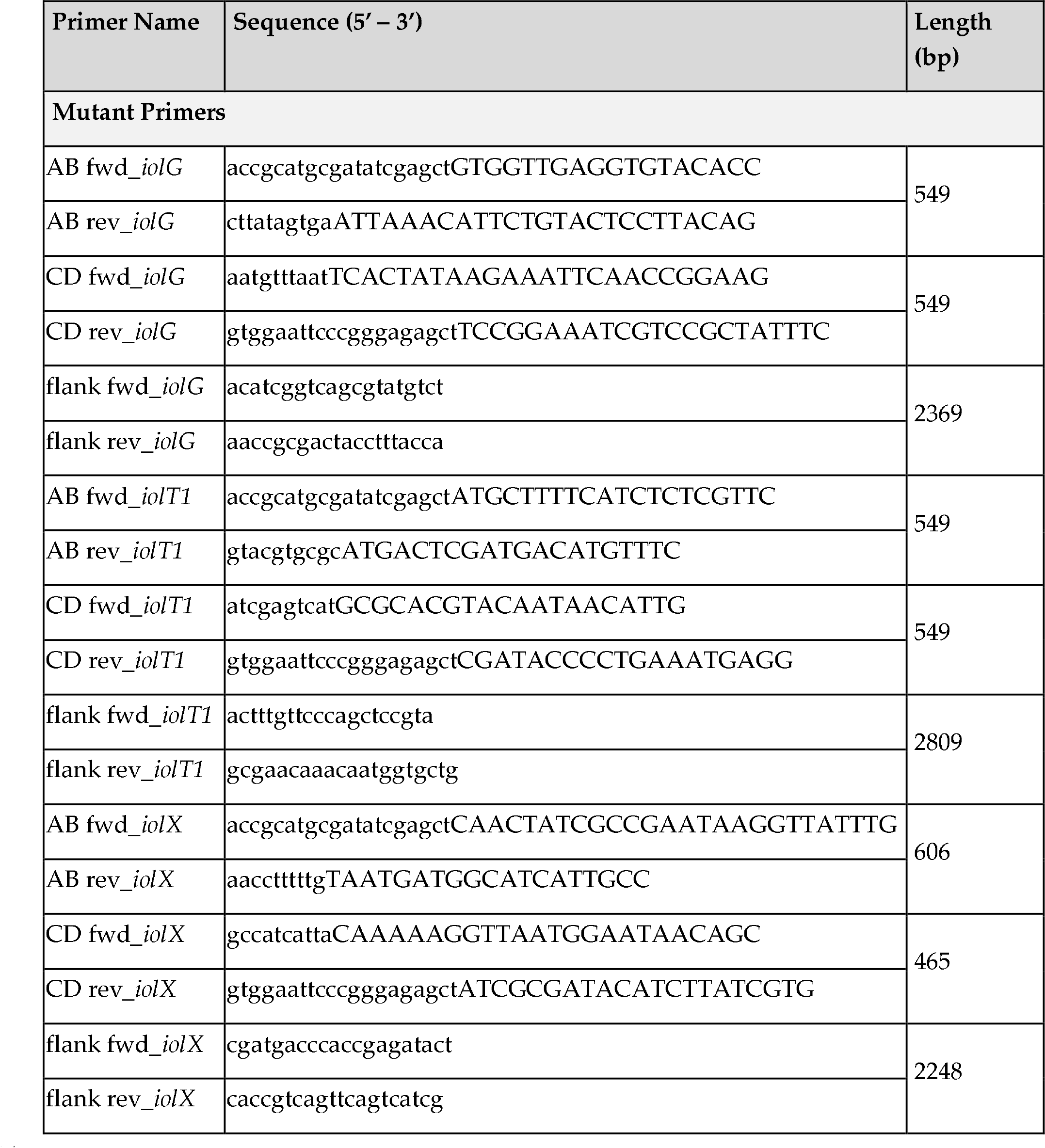
Primers used in this study.

### Metabolic Growth Analysis

Strains were grown overnight in M9G 2% NaCl at 28°C with aeration. Overnight cultures were washed twice with PBS and then inoculated 1:40 into 200 µL fresh M9 media supplemented with 2% NaCl, and a specified carbon source (glucose (20 mM) or *myo*-inositol (20 mM)). Plates were incubated at 28°C with intermittent shaking for 1 min every hour. Optical densities were taken at 595 nm hourly for a total of 30 h using a Tecan Sunrise microplate reader and Magellan plate reader software (Tecan Systems Inc., San Jose, CA).

### Preparation of cellular extracts and proton nuclear magnetic resonance (^1^H-NMR)

Wild type *V. coralliilyticus* was grown overnight at 28°C in M9G 4% NaCl supplemented with 1 mM choline. Stationary phase cells were pelleted and washed twice with PBS. Three freeze-thaw cycles were performed with the cell pellets to increase lysis, and the cells were resuspended in 750 µL ethanol. Debris was pelleted by centrifugation, and the ethanol solution was transferred to a clean tube and evaporated under vacuum. The pellet was resuspended in deuterium oxide (D2O), and insoluble material was removed by centrifugation. The solution was transferred to a 5-mm NMR tube for analysis on a Bruker AVANCE 600NMR spectrometer at a proton frequency of 600.13 MHz with a sweep of 12,376 Hz and a relaxation delay of 5 s. Sixteen scans were co-added for each spectrum.

### Phylogenetic analysis

Phylogenetic analysis was conducted using the program MEGA X (117). The evolutionary history of IolG, IolA and RpoB was inferred using the Neighbor-Joining method (118). The evolutionary distances were computed using the Jones-Taylor-Thornton (JTT) matrix-based method and are in the units of the number of amino acid substitutions per site (85). All ambiguous positions were removed for each sequence pair (pairwise deletion option).

### Ligand Docking Modeling

The transporter amino acid sequences were downloaded from NCBI and aligned using ClustalW(84): *B. subtilis* IolT (<u>WP_003234027.1</u>), *S. enterica* IolT1 (<u>NP_463279.1</u>), *V. coralliilyticus* IolT1 (<u>WP_099607352.1</u>), *V. coralliilyticus* IolT2 (<u>WP_172853823.1</u>), *C. crescentus* IbpA (<u>WP_010918744.1</u>), and *V. coralliilyticus* IolX (<u>WP_227495657.1</u>). Transporter alignments were displayed and annotated using ESPript (http://espript.ibcp.fr/ESPript/cgi-bin/ESPript.cgi) (119). Predicted protein structures were obtained from AlphaFold (120). The Ligand model for *myo*-inositol (MI) was obtained from Chemical Entities of Biological Interest (CheBI) EMBL (121). Polar hydrogens were added to the ligand structures. The docking experiments were performed using SeamDock which was set to use docking parameters for the Vina software (Spacing: 1.0, Mode: 2, Energy Range: 5, Exhaustiveness: 8) (122).

## Data Availability

All datasets are available upon request.

## ACKNOWLEDGEMENTS

This research was supported in part by a National Science Foundation grant (award IOS-1656688) to E.F.B. KBL was funded in part by the Chemistry-Biology Interface predoctoral training program grant: 5T32GM008550.

