## Supplementary data figures S1-S12, Tables S1-S6 for "Osmotic stress response of the coral and oyster pathogen *Vibrio coralliilyticus*: acquisition of catabolism gene clusters for the compatible solute and signaling molecule *myo* -inositol"

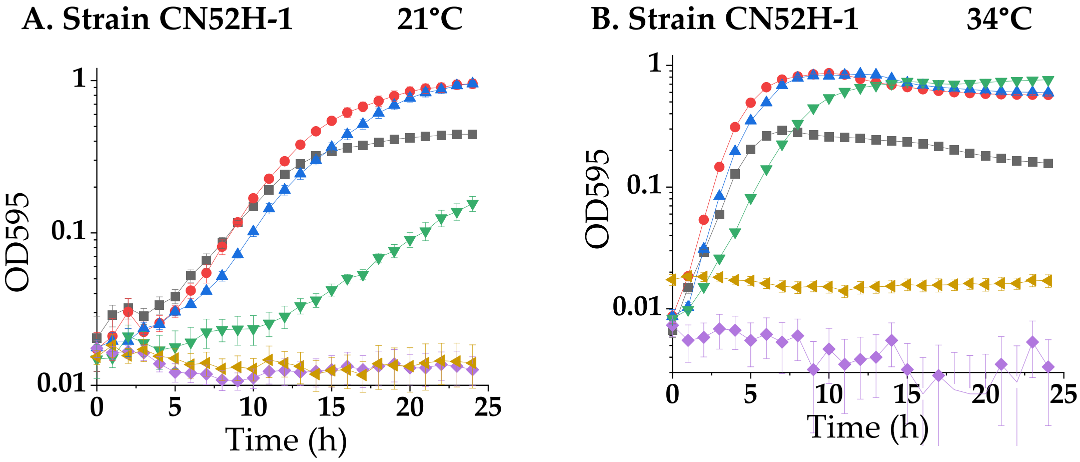

**Fig. S1.** ***Vibrio coralliilyticus*** **growth at different temperatures.** Growth curves of *V. coralliilyticus* CN52H-1, in minimal media with glucose (M9G) containing 1% to 6% NaCl. Growth was measured every hour for 24 h for strain **A.** at 21°C, **B.** at 34°C. Mean and standard deviation of three biological replicates are shown.

**A. H-NMR spectra of *V. coralliilyticus* OCN014 in the absence of choline**

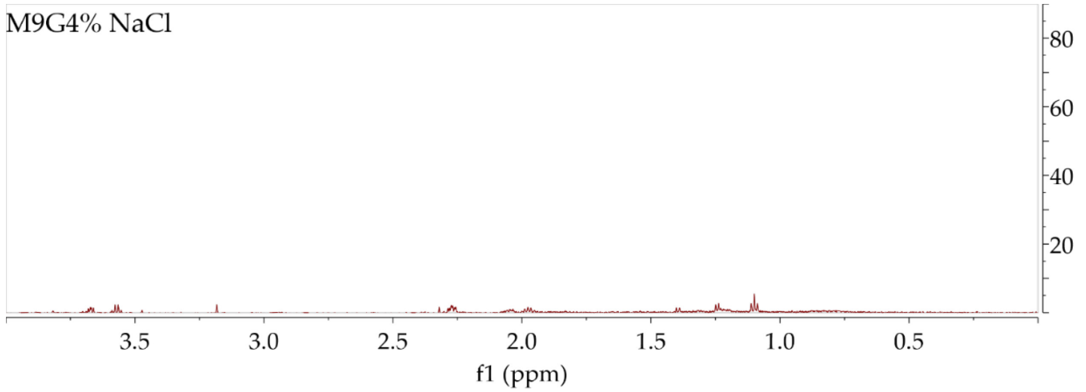

**B. H-NMR spectra of *V. coralliilyticus* OCN014 in the presence of choline**

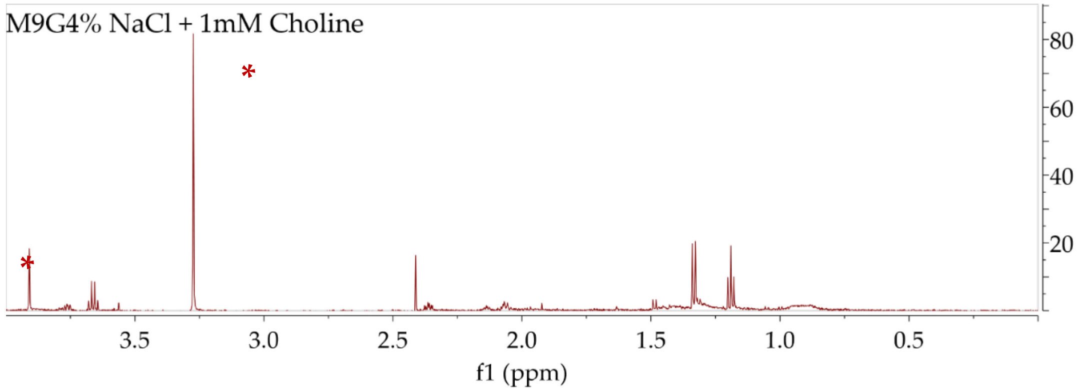

**Fig. S2. *V. coralliilyticus* compatible solute biosynthesis. A.** $^{1}$H-NMR spectra of *V. coralliilyticus* OCN014 cellular extract grown in minimal media (M9G) 4% NaCl without choline and **B.** with the addition of 1 mM choline. The spectral peaks corresponding to glycine betaine are labeled with red asterisks.

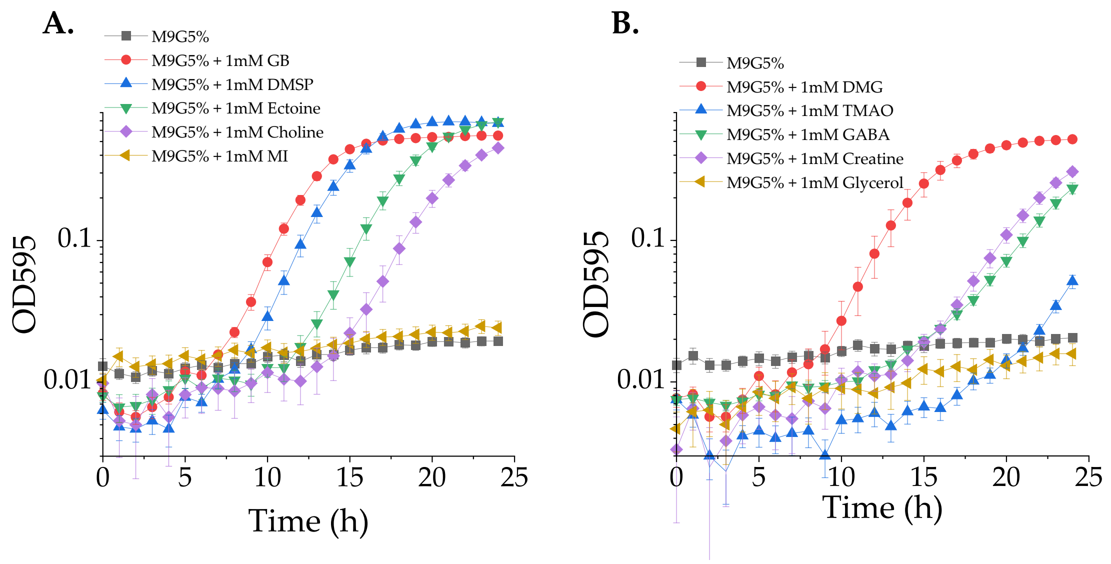

**Fig. S3.** ***V. coralliilyticus*** **can utilize seven of the ten tested compatible solutes as osmoprotectants.** Growth curve analysis of *V. coralliilyticus* in minimal media containing glucose (M9G) and **(A)** 5% NaCl with the addition of 1mM GB (glycine betaine), DMSP (dimethylsulfoniopropionate), Ectoine, Choline, MI (myo-inositol) **(B)** DMG (dimethylglycine), trimethylamine N-oxide (TMAO), gamma-aminobutyric acid (GABA), Creatine, and Glycerol. Growth was measured every hour for 24hr at 28°C**.** Mean and standard deviation of three biological replicates are shown

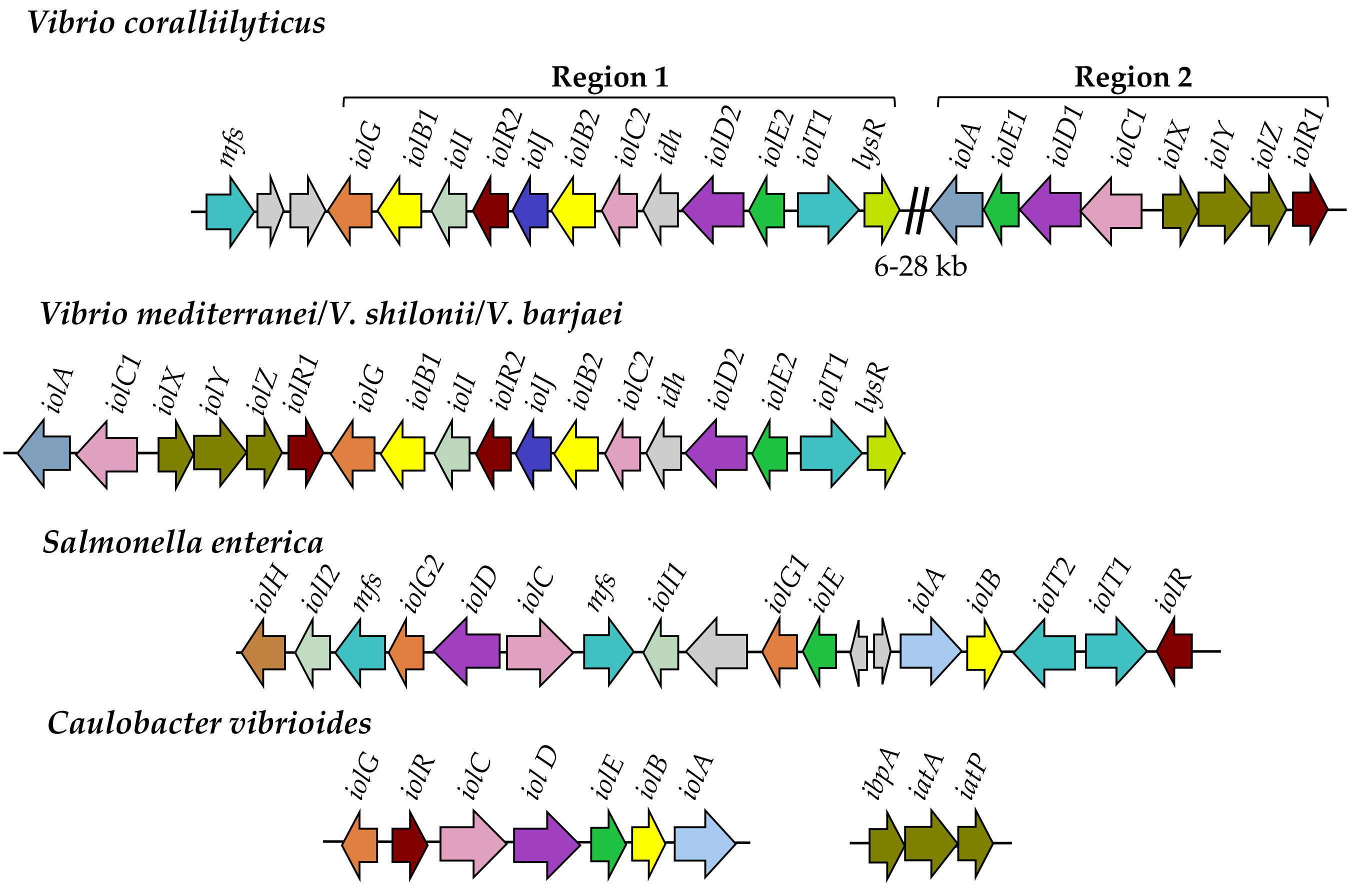

| | |
|---|---|
| **IolA Methylmalonic semialdehyde dehydrogenase** | **IolG Inositol dehydrogenase** |
| **IolB 5-deoxy-glucuronate isomerase** | **IolJ Bisphosphate Aldolase** |
| **IolC 5-dehydro-2 deoxygluconokinase** | IolR MSF transporter |
| **IolD Acylhydrolase** | IolT MSF transporter |
| **IolE Myo-inosose dehydratase** | IolXYZ/Ibp ABC transporter |

**Fig. S4. High variability in chromosomal organization of *iol* genes among bacteria.** Schematics of the gene order of *myo*-inositol transporter, catabolism and regulatory genes among previously characterized species and *Vibrio*. Arrows represent open reading frames and direction represents direction of transcription. Arrows of similar color represent similar proteins/genes.

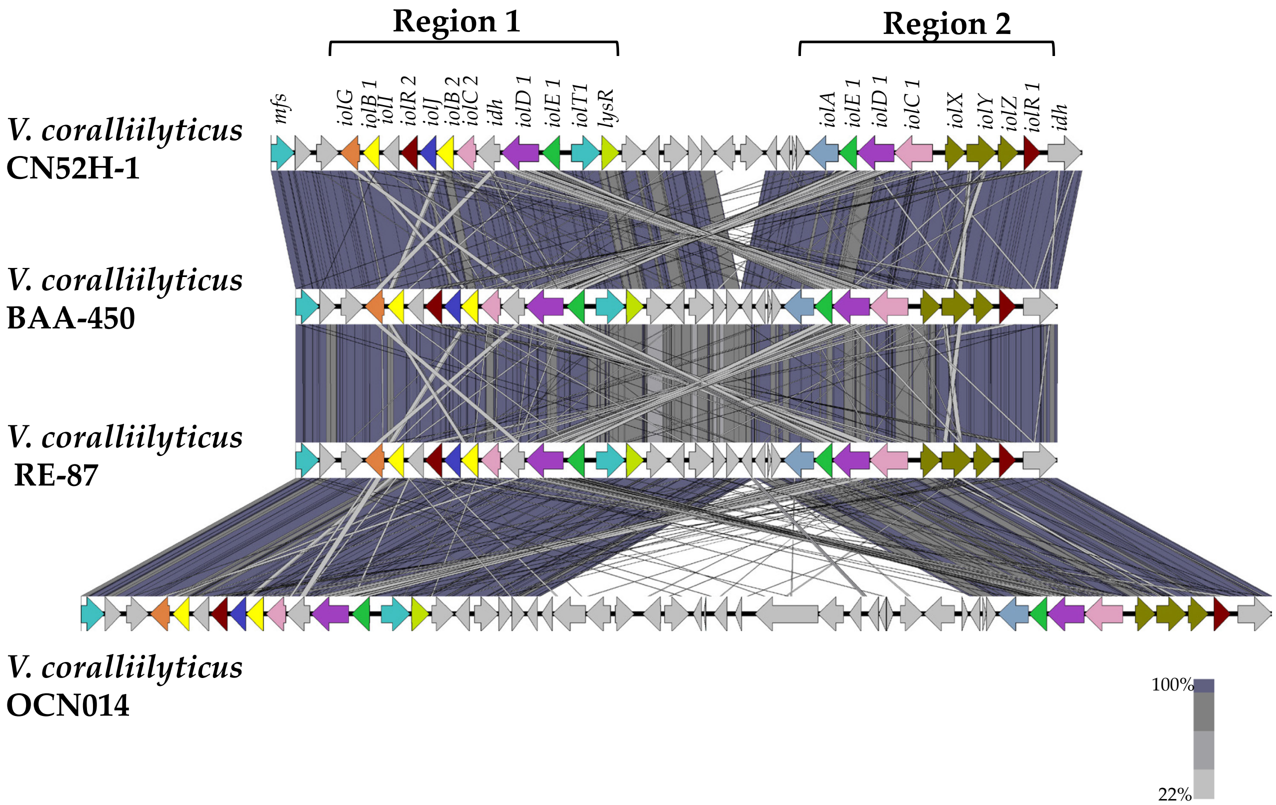

**Fig. S5. Homology of *V. coralliilyticus myo*-inositol gene clusters is conserved in all strains.** Shown here are V. coralliilyticus strains CN52H-1, BAA-450, RE-87, and OCN014. Schematics of the gene order of *myo*-inositol transporter, catabolism and regulatory genes shown. Arrows represent open reading frames and direction represents direction of transcription. Arrows of similar color represent similar proteins/genes.

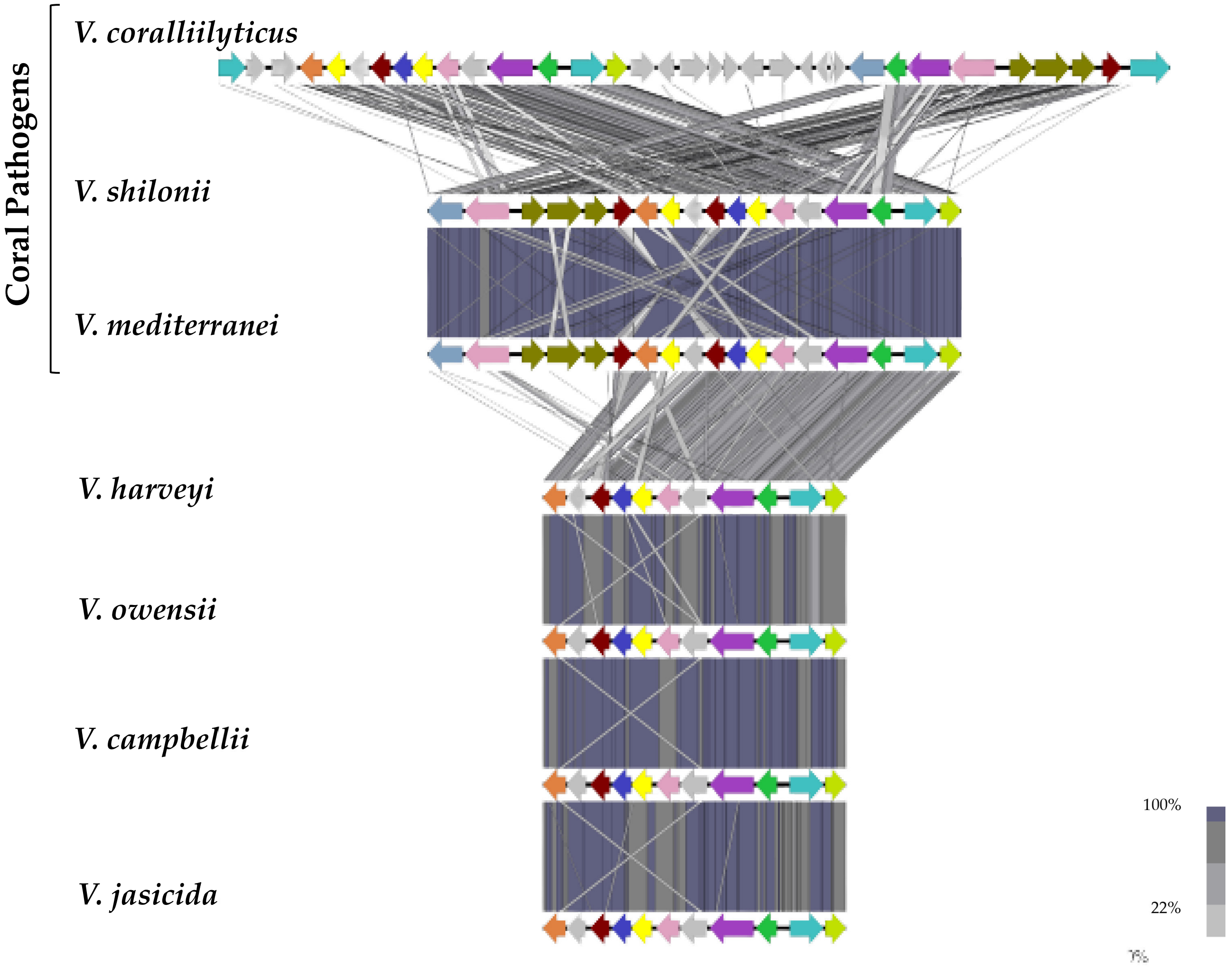

**Fig. S6. Gene homology among *Vibrio* species**. Schematics of the gene order of *myo*-inositol transporter, catabolism and regulatory genes shown. Arrows represent open reading frames and direction represents direction of transcription. Arrows of similar color represent similar proteins/genes.

| ***V. coralliilyticus* IolT1** (WP_099607352.1) | |
|---|---|
| Affinity (kcal/mol) | -6.2 |
| Predicted Binding Residues | Q274 |
| | N279 |
| | Y368 |
| | Q400 |

| ***S. enterica* IolT1** (NP_463279.1) | |
|---|---|
| Affinity (kcal/mol) | -6.1 |
| Predicted Binding Residues | Q269 |
| | Q270 |
| | Y364 |
| | W373 |
| | M396 |

| ***B. subtilis* IolT** (WP_003234027.1) | |
|---|---|
| Affinity (kcal/mol) | -7.0 |
| Predicted Binding Residues | Q267 |
| | Q268 |
| | N273 |
| | Q366 |
| | Q367 |
| | S371 |
| | W375 |

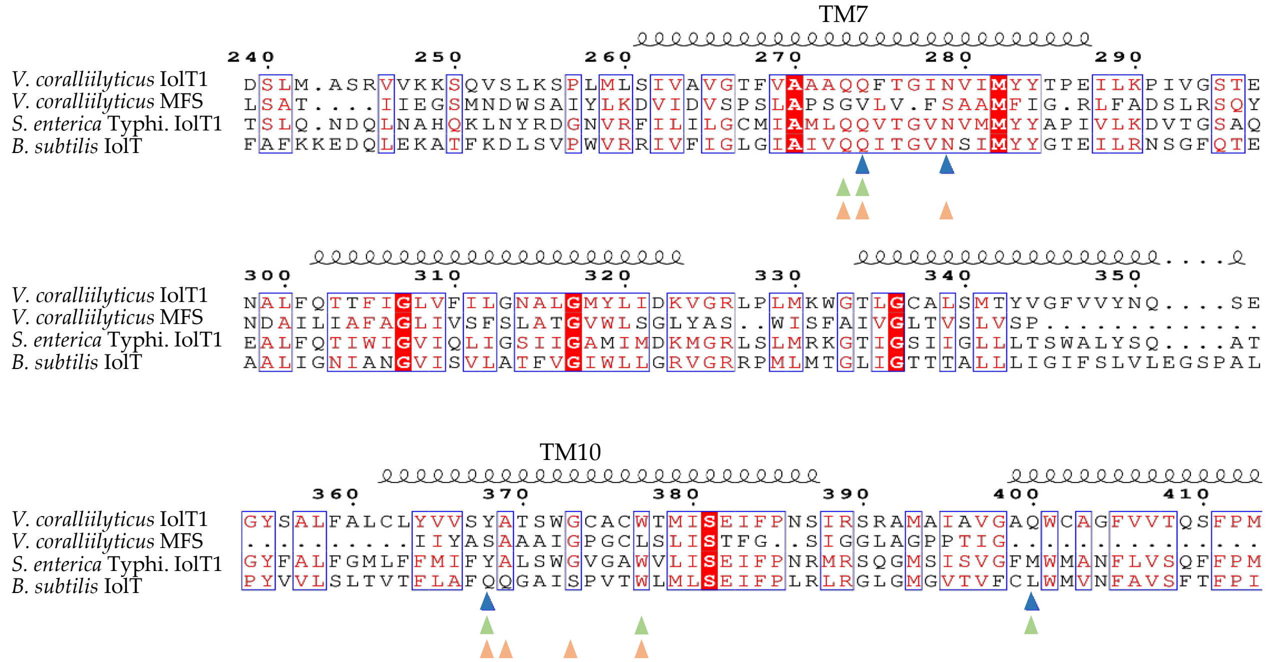

**Fig. S7 . Favorable *myo*-inositol ligand docking predicted for *V. coralliilyticus* MFS-type transporter IolT1.** Results of *myo*-inositol docking study showing free energy change upon substrate binding and putative residues mediating the interactions with the ligand. *B. subtilis* IolT shown to take up *myo*-inositol by Yoshida et al., 2002 and *S. enterica* IolT1 shown to take up *myo*-inositol by Kroger et al., 2010. *V. coralliilyticus* IolT1 has 93% coverage, 34.6% amino acid identity to *B. subtilis* IolT and 99% coverage, 48.8% amino acid identity to *S. enterica* IolT1. Residues that are predicted to interact with *myo*-inositol are shown with an up arrow.

| *V. coralliilyticus* **IolX** (WP_227495657.1) | |
|---|---|
| Affinity (kcal/mol) | -5.3 |
| Putative binding residues | D63 |
| | N141 |
| | Q187 |
| | D269 |
| | Q289 |

| *C. vibrioides* (formerly *crescentus)* **IbpA** (WP_010918744.1) | | |
|---|---|---|
| Affinity (kcal/mol) | -7.4 | Binding Residues: Herrou & Crosson, 2013 |
| Putative binding residues | Q49 | Q49 |
| | D125 | D125 |
| | R126 | R126 |
| | | N168 |
| | D169 | D169 |
| | S174 | S174 |
| | R178 | R178 |
| | | S203 |
| | N231 | N231 |
| | D258 | D258 |
| | Q278 | |

**B. Predicted *myo*-inositol ligand docking in IolX (ABC-type transporter)**

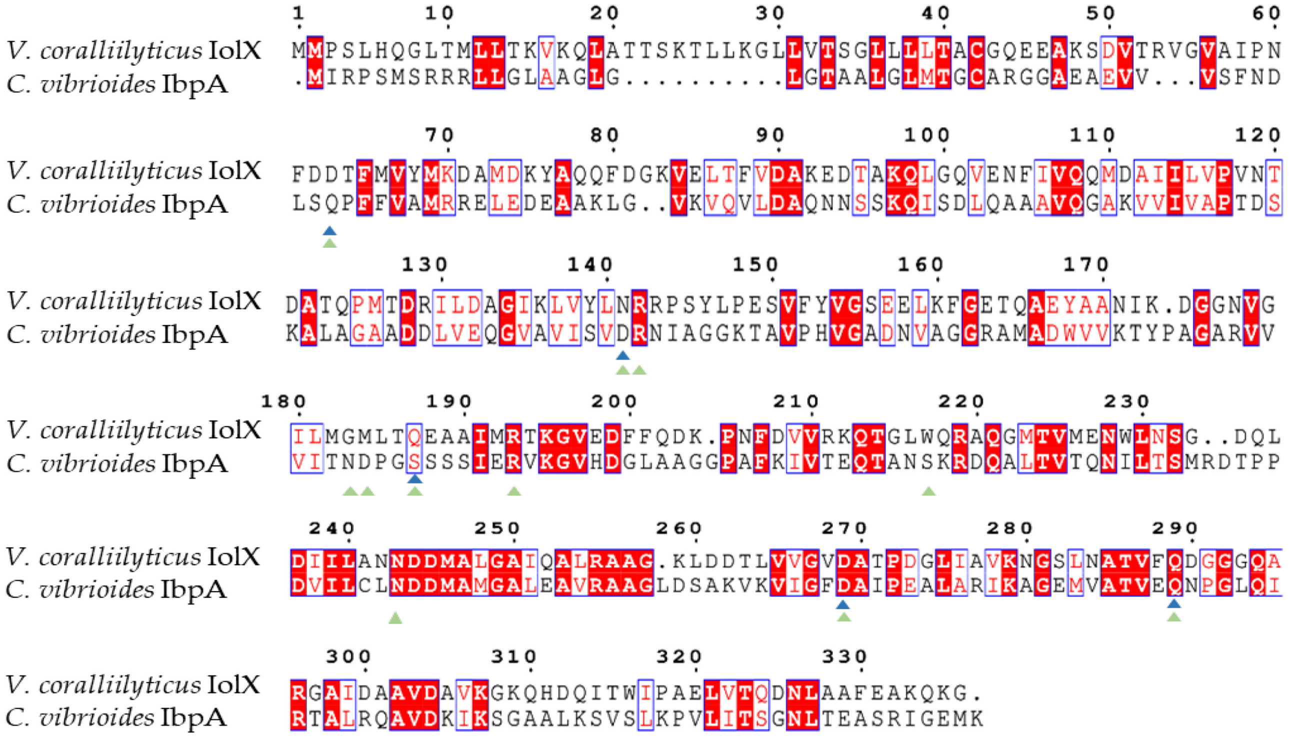

**Fig. S8. Favorable *myo*-inositol ligand docking predicted for *V. coralliilyticus* ABC type transporter IolXYZ.** ABC-type transporter substrate binding proteins; *V. coralliilyticus* IolX (WP_227495657.1, blue arrows) and *C. vibrioides* IbpA (WP_010918744.1, green arrows). *C. vibrioides* has been shown to uptake *myo*-inositol using this ABC-type transporter.

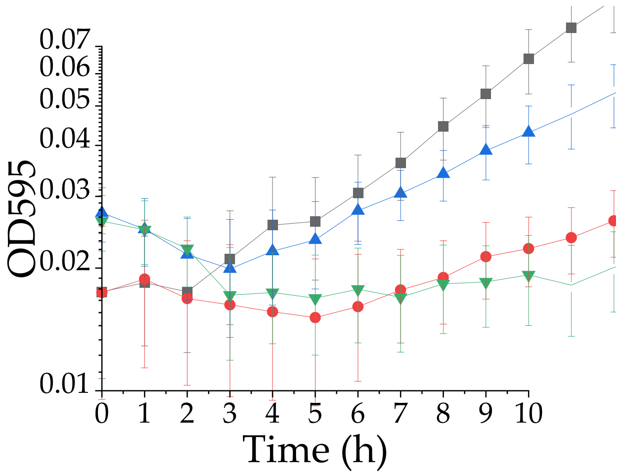

**Fig. S9.** Magnification of growth analysis of wild type (black), Δ*iolT1* (red), Δ*iolX* (blue), and double mutant Δ*iolT1*/Δ*iolX* (green) in minimal media (M9) 2% NaCl with 20 mM *myo*-inositol as a sole carbon source from Fig. 5B. This more clearly shows lag phase differences among mutant strains and wild type.

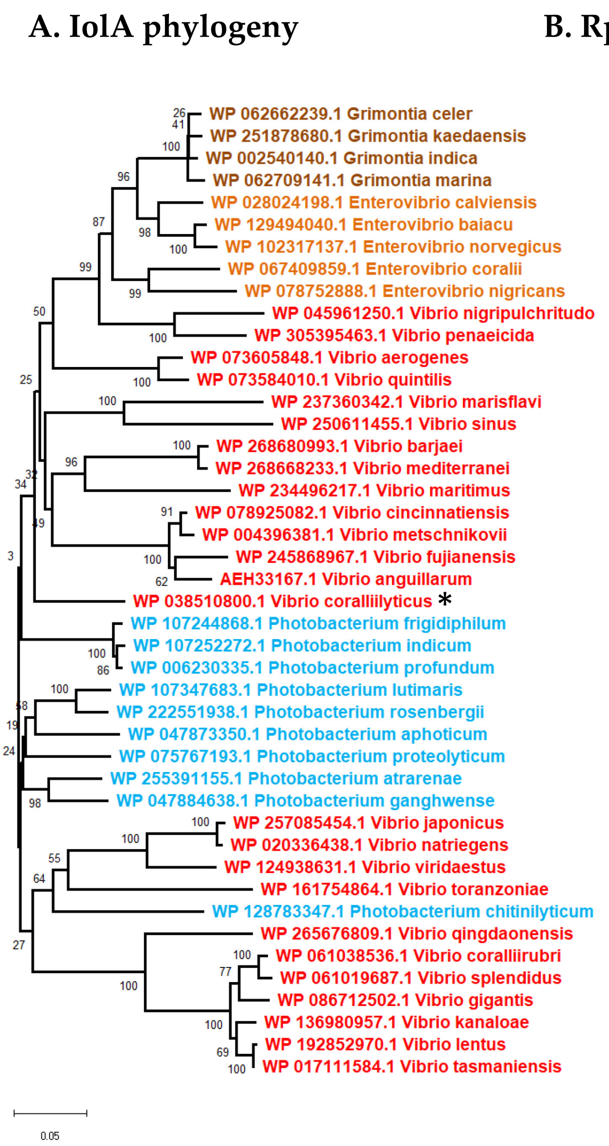

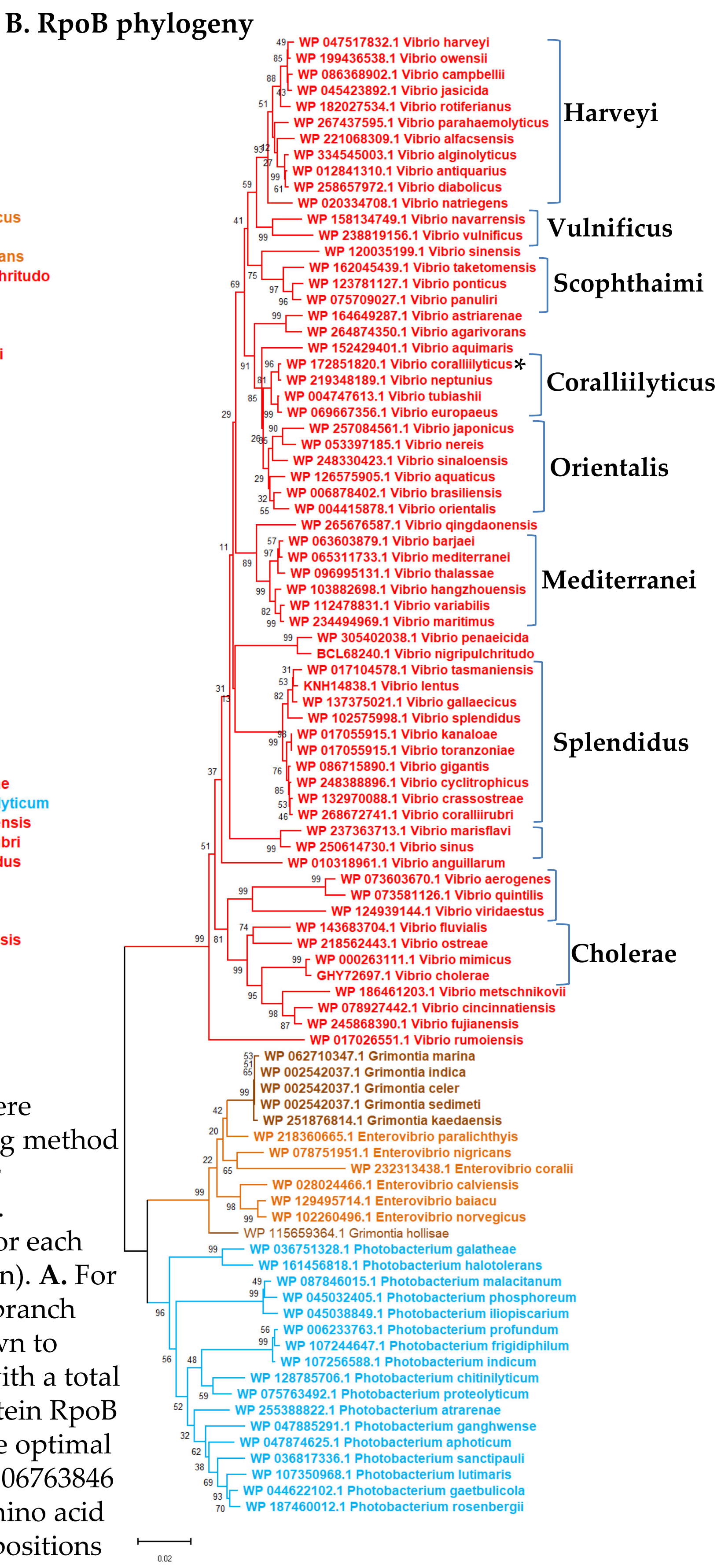

**Fig. S10. Phylogeny of IolA and the housekeeping protein RpoB**. Trees were constructed using the Neighbor-Joining method with the JTT matrix-based method for distances with bootstrap value of 1000. Ambiguous positions were removed for each sequence pair (pairwise deletion option). **A.** For IolA the optimal tree with the sum of branch length = 2.37747408 is shown and drawn to scale from 44 amino acid sequences, with a total of 510 positions. **B**. Housekeeping protein RpoB showing the major clades of *Vibrio*. The optimal tree with the sum of branch length = 1.06763846 is shown. This analysis involved 92 amino acid sequences. There were a total of 1344 positions in the final dataset.

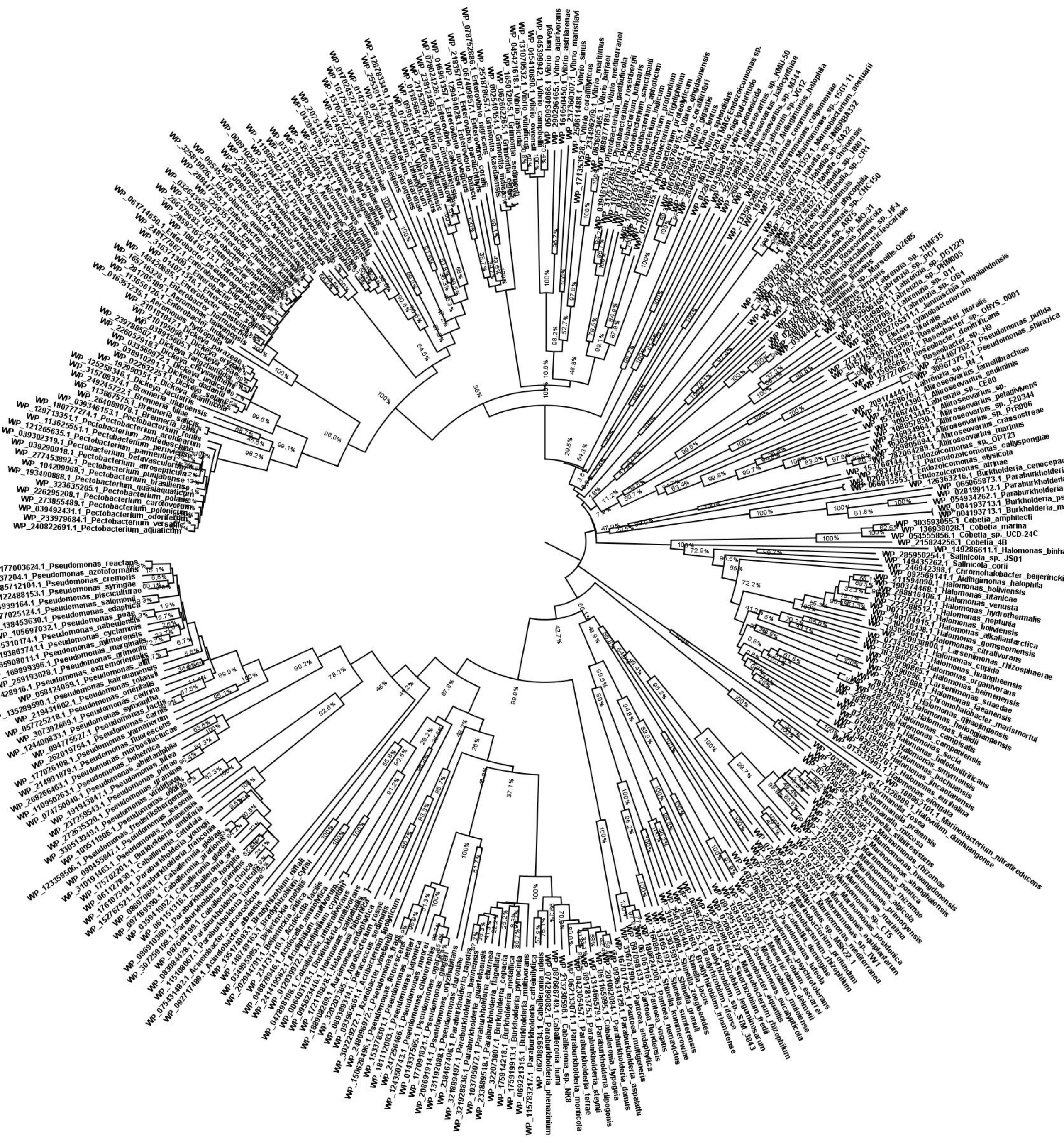

**Fig. S11. Evolutionary analysis of IolG among marine bacteria**. The optimal tree is shown. The tree inferred using the Neighbor-Joining method is drawn to scale, with branch lengths in the same units as those of the evolutionary distances used to infer the phylogenetic tree. The evolutionary distances were computed using the p-distance method and are in the units of the number of amino acid differences per site. The rate variation among sites was modeled with a gamma distribution (shape parameter = 3). This analysis involved 358 amino acid sequences. All ambiguous positions were removed for each sequence pair (pairwise deletion option). There were a total of 359 positions in the final dataset. Evolutionary analyses were conducted in MEGA11.

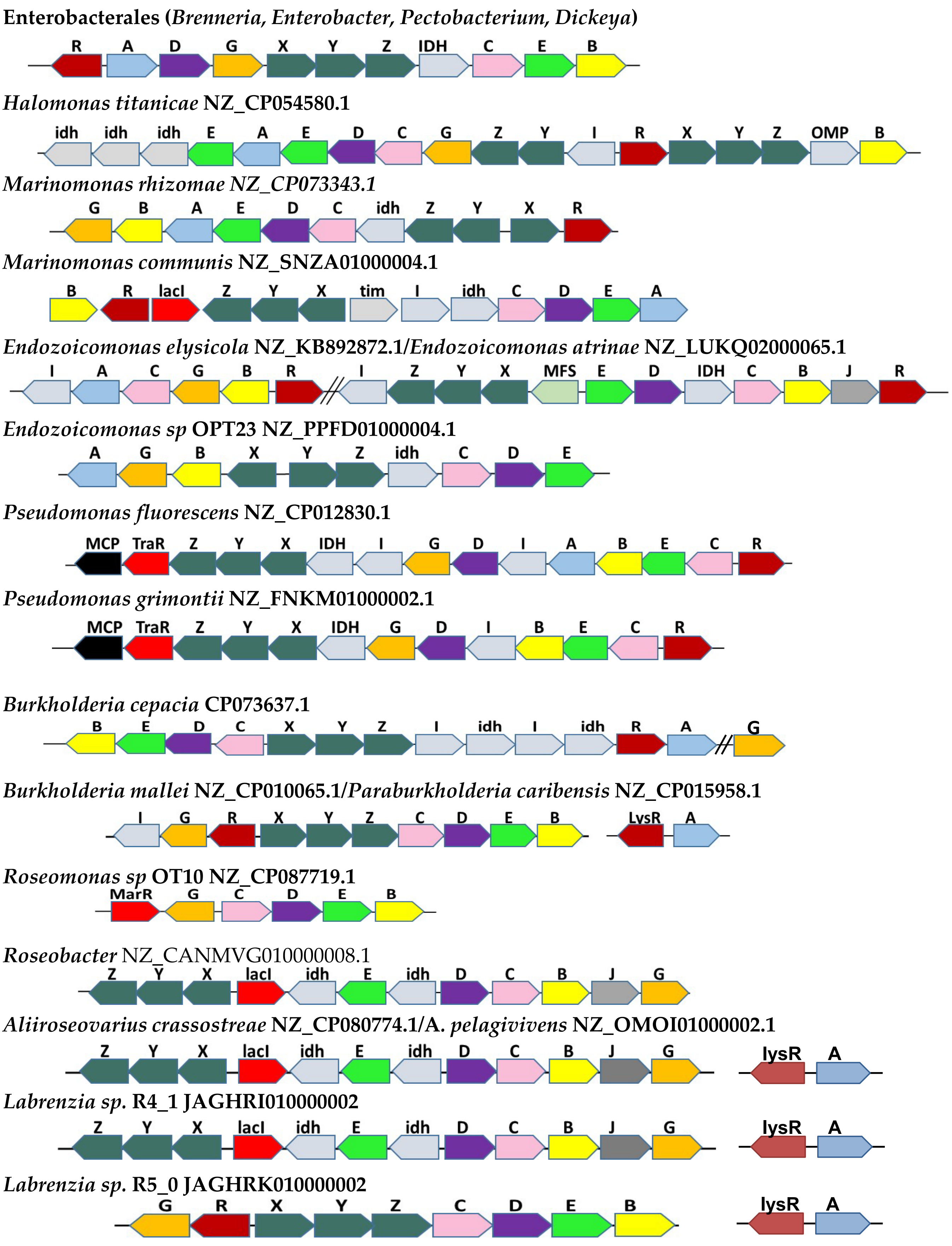

**Fig. S12. Iol cluster organization among marine species.** Arrows represent open reading frames and direction of transcription.

**Table S1.** ***Vibrio coralliilyticus iol*** **gene homology to** ***iol*** **genes in** ***Salmonella enterica***

| *V. coralliilyticus* CN52H-1 | Locus tag | Protein ID | Product | Homology *S. enterica* |
|---|---|---|---|---|
| ***iolE2*** | HRJ43_RS06580 | WP_006962651.1 | myo-inosose-2 dehydratase | 38% |
| ***iolD2*** | HRJ43_RS06575 | WP_172853826.1 | 3D-(3,5/4)-trihydroxycyclohexane-1,2-dione acylhydrolase | 38% |
| ***idh*** | HRJ43_RS06570 | WP_043008502.1 | D-lactate dehydrogenase | none |
| ***iolC2*** | HRJ43_RS06565 | WP_019274787.1 | 5-dehydro-2-deoxygluconokinase | 33% |
| ***iolB2*** | HRJ43_RS06560 | WP_006962643.1 | 5-deoxy-glucuronate isomerase | 31% |
| ***iolJ*** | HRJ43_RS06555 | WP_006962641.1 | class II fructose-bisphosphate aldolase | none |
| ***iolR2*** | HRJ43_RS06550 | WP_006962640.1 | MurR/RpiR family regulator | 40% |
| ***iolI*** | HRJ43_RS06545 | WP_172853825.1 | sugar phosphate isomerase/epimerase | 21% |
| ***iolB1*** | HRJ43_RS06540 | WP_172853824.1 | 5-deoxy-glucuronate isomerase | 64% |
| ***iolG*** | HRJ43_RS06535 | WP_038162996.1 | inositol dehydrogenase | 36% |
| ***iolT1*** | HRJ43_RS06585 | WP_099607352.1 | MFS transporter | 49% |
| ***lysR*** | HRJ43_RS06590 | WP_172853827.1 | transcriptional regulator | none |
| ***iolC1*** | HRJ43_RS06665 | WP_099607365.1 | 5-dehydro-2-deoxygluconokinase | 67% |
| ***iolD1*** | HRJ43_RS06660 | WP_172853836.1 | 3D-(3,5/4)-trihydroxycyclohexane-1,2-dione acylhydrolase | 44% |
| ***iolE1*** | HRJ43_RS06655 | WP_172853835.1 | myo-inosose-2 dehydratase | 38% |
| ***iolA*** | HRJ43_RS06650 | WP_172853834.1 | CoA-acylating methylmalonate-semialdehyde dehydrogenase | 65% |
| ***iolX*** | HRJ43_RS06670 | WP_227495657.1 | substrate-binding domain-containing protein | none |
| ***iolY*** | HRJ43_RS06675 | WP_006962669.1 | sugar ABC transporter ATP-binding protein | none |
| ***iolZ*** | HRJ43_RS06680 | WP_006962670.1 | ABC transporter permease | none |
| ***iolR1*** | HRJ43_RS06685 | WP_006962671.1 | MurR/RpiR family regulator | 67% |

**Table S2.** ***Pectobacterium brasiliense iol*** **cluster genome location (tRNA-Asn), associated mobile genetic element (in brown and integases in red) and addiction system (in green).**

| Genome location | Strand | Protein | Gene | Locus tag | Protein tag | mino aci |
|---|---|---|---|---|---|---|
| NZ_CP024780.1:1631671-1631746 | plus | **tRNA-Asn** | | CTV95_RS07455 | | |
| NZ_CP024780.1:1631908-1633179 | plus | **integrase arm-type DNA-binding domain-containing protein** | | CTV95_RS07460 | 1 WP_100017225.1 | 423 |
| NZ_CP024780.1:1633313-1633621 | minus | helix-turn-helix transcriptional regulator | | CTV95_RS07465 | 1 WP_318841327.1 | 102 |
| NZ_CP024780.1:1633813-1634127 | minus | helix-turn-helix transcriptional regulator | | CTV95_RS07470 | 1 WP_014699384.1 | 104 |
| NZ_CP024780.1:1634274-1634546 | plus | helix-turn-helix transcriptional regulator | | CTV95_RS07475 | 1 WP_014699383.1 | 90 |
| NZ_CP024780.1:1634962-1635876 | plus | hypothetical protein | | CTV95_RS07480 | 1 WP_100017227.1 | 304 |
| NZ_CP024780.1:1636030-1636224 | plus | AlpA family transcriptional regulator | | CTV95_RS07485 | 1 WP_039359985.1 | 64 |
| NZ_CP024780.1:1636241-1636825 | plus | DUF2857 domain-containing protein | | CTV95_RS07490 | 1 WP_100017230.1 | 194 |
| NZ_CP024780.1:1636815-1637024 | plus | TraR/DksA family transcriptional regulator | | CTV95_RS07495 | 1 WP_015730135.1 | 69 |
| NZ_CP024780.1:1637352-1638236 | minus | hypothetical protein | | CTV95_RS07500 | 1 WP_100017232.1 | 294 |
| NZ_CP024780.1:1638664-1639449 | minus | hypothetical protein | | CTV95_RS22560 | 1 WP_198406574.1 | 261 |
| NZ_CP024780.1:1640192-1640413 | plus | hypothetical protein | | CTV95_RS07510 | 1 WP_100017234.1 | 73 |
| NZ_CP024780.1:1640495-1641160 | plus | lytic transglycosylase domain-containing protein | | CTV95_RS07515 | 1 WP_100017236.1 | 221 |
| NZ_CP024780.1:1641157-1641438 | plus | TrbC/VirB2 family protein | | CTV95_RS07520 | 1 WP_100017238.1 | 93 |
| NZ_CP024780.1:1641448-1644195 | plus | VirB3 family type IV secretion system protein | | CTV95_RS07525 | 1 WP_100017239.1 | 915 |
| NZ_CP024780.1:1644219-1644935 | plus | type IV secretion system protein | | CTV95_RS07530 | 1 WP_100017241.1 | 238 |
| NZ_CP024780.1:1644946-1645170 | plus | EexN family lipoprotein | | CTV95_RS07535 | 1 WP_100017243.1 | 74 |
| NZ_CP024780.1:1645182-1646222 | plus | type IV secretion system protein | | CTV95_RS07540 | 1 WP_100017245.1 | 346 |
| NZ_CP024780.1:1646310-1646450 | plus | hypothetical protein | | CTV95_RS22565 | 1 WP_181847862.1 | 46 |
| NZ_CP024780.1:1646443-1647126 | plus | type IV secretion system protein | | CTV95_RS07545 | 1 WP_100017246.1 | 227 |
| NZ_CP024780.1:1647123-1648016 | plus | P-type conjugative transfer protein VirB9 | virB9 | CTV95_RS07550 | 1 WP_100017248.1 | 297 |
| NZ_CP024780.1:1648013-1649245 | plus | VirB10/TraB/TrbI family type IV secretion system protein | virB10 | CTV95_RS07555 | 1 WP_100017250.1 | 410 |
| NZ_CP024780.1:1649235-1650251 | plus | P-type DNA transfer ATPase VirB11 | virB11 | CTV95_RS07560 | 1 WP_100017252.1 | 338 |
| NZ_CP024780.1:1650294-1650596 | plus | TrbM/KikA/MpfK family conjugal transfer protein | | CTV95_RS07565 | 1 WP_100017254.1 | 100 |
| NZ_CP024780.1:1651317-1653179 | plus | type IV secretory system conjugative DNA transfer family protein | | CTV95_RS07575 | 1 WP_100017255.1 | 620 |
| NZ_CP024780.1:1653188-1653934 | plus | MobC family replication-relaxation protein | mobC | CTV95_RS07580 | 1 WP_100017257.1 | 248 |
| NZ_CP024780.1:1654160-1654714 | plus | hypothetical protein | | CTV95_RS07585 | 1 WP_198406575.1 | 184 |
| NZ_CP024780.1:1654963-1655904 | minus | hypothetical protein | | CTV95_RS07590 | 1 WP_100017258.1 | 313 |
| NZ_CP024780.1:1656240-1657007 | minus | hypothetical protein | | CTV95_RS07595 | 1 WP_226302438.1 | 255 |
| NZ_CP024780.1:1657675-1658607 | plus | hypothetical protein | | CTV95_RS07600 | 1 WP_100017260.1 | 310 |
| NZ_CP024780.1:1658884-1661178 | plus | AAA family ATPase | | CTV95_RS07605 | 1 WP_100017262.1 | 764 |
| NZ_CP024780.1:1661175-1663022 | plus | UvrD-helicase domain-containing protein | | CTV95_RS07610 | 1 WP_100017264.1 | 615 |
| NZ_CP024780.1:1663371-1664303 | minus | S-4TM family putative pore-forming effector | | CTV95_RS07615 | 1 WP_100017266.1 | 310 |
| NZ_CP024780.1:1664307-1665302 | minus | nucleotidyltransferase | | CTV95_RS07620 | 1 WP_100017268.1 | 331 |
| NZ_CP024780.1:1665452-1666351 | minus | SMEK domain-containing protein | | CTV95_RS07625 | 1 WP_100017269.1 | 299 |
| NZ_CP024780.1:1666629-1666763 | plus | uncharacterized gene | | CTV95_RS22865 | | |
| NZ_CP024780.1:1666797-1667111 | minus | CcdB family protein | | CTV95_RS07635 | 1 WP_100017271.1 | 104 |
| NZ_CP024780.1:1667114-1667350 | minus | type II toxin-antitoxin system CcdA family antitoxin | | CTV95_RS07640 | 1 WP_100017272.1 | 78 |
| NZ_CP024780.1:1667670-1668614 | minus | zincin-like metallopeptidase domain-containing protein | | CTV95_RS07645 | 1 WP_100017273.1 | 314 |
| NZ_CP024780.1:1669095-1669391 | plus | DUF3892 domain-containing protein | | CTV95_RS07650 | 1 WP_100017274.1 | 98 |
| NZ_CP024780.1:1669596-1670198 | minus | **site-specific integrase** | | CTV95_RS07655 | 1 WP_065684227.1 | 200 |
| NZ_CP024780.1:1670383-1671969 | minus | hypothetical protein | | CTV95_RS07660 | 1 WP_157765229.1 | 528 |
| NZ_CP024780.1:1672045-1673340 | minus | FRG domain-containing protein | | CTV95_RS07665 | 1 WP_180745091.1 | 431 |
| NZ_CP024780.1:1674099-1674263 | minus | uncharacterized gene | | CTV95_RS22725 | | |
| NZ_CP024780.1:1674416-1675054 | minus | NAD(P)H-binding protein | | CTV95_RS07675 | 1 WP_100017276.1 | 212 |
| NZ_CP024780.1:1675300-1676199 | plus | LysR family transcriptional regulator | | CTV95_RS07680 | 1 WP_100017277.1 | 299 |
| NZ_CP024780.1:1676599-1677513 | plus | LysR family transcriptional regulator | | CTV95_RS07690 | 1 WP_100017278.1 | 304 |
| NZ_CP024780.1:1677539-1678129 | minus | flavin reductase family protein | | CTV95_RS07695 | 1 WP_100017279.1 | 196 |
| NZ_CP024780.1:1678512-1678844 | plus | DUF1971 domain-containing protein | | CTV95_RS07700 | 1 WP_100017280.1 | 110 |
| NZ_CP024780.1:1678855-1679193 | plus | DUF1869 domain-containing protein | | CTV95_RS07705 | 1 WP_100017281.1 | 112 |
| NZ_CP024780.1:1679415-1680479 | minus | sensor domain-containing diguanylate cyclase | | CTV95_RS07710 | 1 WP_100017282.1 | 354 |
| NZ_CP024780.1:1680921-1682786 | plus | beta-glucoside-specific PTS transporter subunit IIABC | | CTV95_RS07715 | 1 WP_100017283.1 | 621 |
| NZ_CP024780.1:1682786-1684249 | plus | family 1 glycosylhydrolase | | CTV95_RS07720 | 1 WP_039468283.1 | 487 |
| NZ_CP024780.1:1684343-1685182 | plus | PRD domain-containing protein | | CTV95_RS07725 | 1 WP_039499196.1 | 279 |
| NZ_CP024780.1:1685416-1686465 | plus | Fe(3+) ABC transporter substrate-binding protein | | CTV95_RS07730 | 1 WP_100017284.1 | 349 |
| NZ_CP024780.1:1686449-1688146 | plus | iron ABC transporter permease | | CTV95_RS07735 | 1 WP_100017285.1 | 565 |
| NZ_CP024780.1:1688168-1688857 | plus | ABC transporter ATP-binding protein | | CTV95_RS07740 | 1 WP_100017286.1 | 229 |
| NZ_CP024780.1:1688935-1689213 | plus | type II toxin-antitoxin system ParD family antitoxin | | CTV95_RS07745 | 1 WP_100017287.1 | 92 |
| NZ_CP024780.1:1689213-1689533 | plus | type II toxin-antitoxin system RelE/ParE family toxin | | CTV95_RS07750 | 1 WP_014914766.1 | 106 |
| NZ_CP024780.1:1689636-1690472 | minus | 5-deoxy-glucuronate isomerase | **iolB** | CTV95_RS07755 | 1 WP_039554464.1 | 278 |
| NZ_CP024780.1:1690561-1691451 | minus | myo-inosose-2 dehydratase | **iolE** | CTV95_RS07760 | 1 WP_014914764.1 | 296 |
| NZ_CP024780.1:1691485-1693389 | minus | 5-dehydro-2-deoxygluconokinase | **iolC** | CTV95_RS07765 | 1 WP_010281444.1 | 634 |
| NZ_CP024780.1:1693584-1694717 | minus | Gfo/Idh/MocA family oxidoreductase | **idh** | CTV95_RS07770 | 1 WP_039287661.1 | 377 |
| NZ_CP024780.1:1694758-1695789 | minus | ABC transporter permease | **iolZ** | CTV95_RS07775 | 1 WP_010281439.1 | 343 |
| NZ_CP024780.1:1695802-1697349 | minus | sugar ABC transporter ATP-binding protein | **iolY** | CTV95_RS07780 | 1 WP_014914761.1 | 515 |
| NZ_CP024780.1:1697422-1698360 | minus | substrate-binding domain-containing protein | **iolX** | CTV95_RS07785 | 1 WP_010308444.1 | 312 |
| NZ_CP024780.1:1698441-1699427 | minus | inositol 2-dehydrogenase | **iolG** | CTV95_RS07790 | 1 WP_039280085.1 | 328 |
| NZ_CP024780.1:1700144-1702075 | minus | 3D-(3,5/4)-trihydroxycyclohexane-1,2-dione acylhydrolase (decyclizing) | **iolD** | CTV95_RS07795 | 1 WP_100017288.1 | 643 |
| NZ_CP024780.1:1702433-1703944 | minus | CoA-acylating methylmalonate-semialdehyde dehydrogenase | **iolA** | CTV95_RS07800 | 1 WP_014914754.1 | 503 |
| NZ_CP024780.1:1704272-1705129 | plus | MurR/RpiR family transcriptional regulator | **iolR** | CTV95_RS07805 | 1 WP_010281425.1 | 285 |
| NZ_CP024780.1:1705177-1705482 | plus | putative quinol monooxygenase | | CTV95_RS07810 | 1 WP_039287697.1 | 101 |
| NZ_CP024780.1:1705809-1706912 | plus | extracellular solute-binding protein | | CTV95_RS07815 | 1 WP_100017289.1 | 367 |
| NZ_CP024780.1:1707063-1708124 | minus | diguanylate cyclase AdrA | adrA | CTV95_RS07820 | 1 WP_014914750.1 | 353 |
| NZ_CP024780.1:1708357-1709193 | minus | PRD domain-containing protein | | CTV95_RS07825 | 1 WP_014914749.1 | 278 |
| NZ_CP024780.1:1709284-1710711 | minus | glycoside hydrolase family 1 protein | | CTV95_RS07830 | 1 WP_039509309.1 | 475 |
| NZ_CP024780.1:1711055-1712683 | minus | S53 family peptidase | | CTV95_RS07835 | 1 WP_100017290.1 | 542 |
| NZ_CP024780.1:1712702-1713259 | minus | chorismate mutase | | CTV95_RS07840 | 1 WP_039287719.1 | 185 |
| NZ_CP024780.1:1713647-1715788 | minus | YjbH domain-containing protein | | CTV95_RS07845 | 1 WP_100017291.1 | 713 |
| NZ_CP024780.1:1715785-1716567 | minus | capsule biosynthesis GfcC family protein | | CTV95_RS07850 | 1 WP_100017292.1 | 260 |
| NZ_CP024780.1:1716577-1717254 | minus | YjbF family lipoprotein | | CTV95_RS07855 | 1 WP_180791592.1 | 225 |
| NZ_CP024780.1:1717330-1717623 | minus | hypothetical protein | | CTV95_RS07860 | 1 WP_100017294.1 | 97 |
| NZ_CP024780.1:1718072-1719478 | minus | **NADP-dependent phosphogluconate dehydrogenase** | **gndA** | CTV95_RS07865 | 1 WP_010281382.1 | 468 |

**Table S3.** ***Pectobacterium carotovorum iol*** **cluster genome location (tRNA-Asn) and addiction system (in green).**

| Genome location | Strand | Protein | Gene | Locus tag | Protein tag | mino acids |
|---|---|---|---|---|---|---|
| CP051652.1:3215076-3215151 | plus | **tRNA-Asn** | | HER17_14335 | | |
| CP051652.1:3215427-3216446 | minus | **NADP-dependent oxidoreductase** | | HER17_14340 | 1 QLL94049. | 339 |
| CP051652.1:3216454-3217272 | minus | **oxidoreductase** | | HER17_14345 | 1 QLL94050. | 272 |
| CP051652.1:3217367-3218149 | minus | **SDR family oxidoreductase** | | HER17_14350 | 1 QLL94051. | 260 |
| CP051652.1:3218225-3219316 | minus | alkene reductase | | HER17_14355 | 1 QLL94052. | 363 |
| CP051652.1:3219454-3220023 | plus | TetR/AcrR family transcriptional regulator | | HER17_14360 | 1 QLL94053. | 189 |
| CP051652.1:3220102-3221037 | minus | alpha/beta hydrolase | | HER17_14365 | 1 QLL94054. | 311 |
| CP051652.1:3221163-3222077 | plus | LysR family transcriptional regulator | | HER17_14370 | 1 QLL94055. | 304 |
| CP051652.1:3222282-3222872 | minus | flavin reductase family protein | | HER17_14375 | 1 QLL94056. | 196 |
| CP051652.1:3223254-3223586 | plus | DUF1971 domain-containing protein | | HER17_14380 | 1 QLL94057. | 110 |
| CP051652.1:3223597-3223941 | plus | DUF1869 domain-containing protein | | HER17_14385 | 1 QLL94058. | 114 |
| CP051652.1:3224125-3225189 | minus | sensor domain-containing diguanylate cyclase | | HER17_14390 | 1 QLL94059. | 354 |
| CP051652.1:3225631-3227496 | plus | PTS transporter subunit EIIC | | HER17_14395 | 1 QLL94060. | 621 |
| CP051652.1:3227496-3228959 | plus | family 1 glycosylhydrolase | | HER17_14400 | 1 QLL94061. | 487 |
| CP051652.1:3229054-3229893 | plus | PRD domain-containing protein | | HER17_14405 | 1 QLL94062. | 279 |
| CP051652.1:3230128-3231177 | plus | Fe(3+) ABC transporter substrate-binding protein | | HER17_14410 | 1 QLL94063. | 349 |
| CP051652.1:3231161-3232858 | plus | iron ABC transporter permease | | HER17_14415 | 1 QLL94064. | 565 |
| CP051652.1:3232880-3233569 | plus | ABC transporter ATP-binding protein | | HER17_14420 | 1 QLL94065. | 229 |
| CP051652.1:3233647-3233925 | plus | type II toxin-antitoxin system ParD family antitoxin | | HER17_14425 | 1 QLL94066. | 92 |
| CP051652.1:3233925-3234245 | plus | type II toxin-antitoxin system RelE/ParE family toxin | | HER17_14430 | 1 QLL94067. | 106 |
| CP051652.1:3234348-3235184 | **minus** | 5-deoxy-glucuronate isomerase | **iolB** | HER17_14435 | 1 QLL94068. | 278 |
| CP051652.1:3235271-3236161 | **minus** | myo-inosose-2 dehydratase | **iolE** | HER17_14440 | 1 QLL94069. | 296 |
| CP051652.1:3236390-3238294 | **minus** | 5-dehydro-2-deoxygluconokinase | **iolC** | HER17_14445 | 1 QLL94070. | 634 |
| CP051652.1:3238339-3239472 | **minus** | **Gfo/Idh/MocA family oxidoreductase** | **idh** | HER17_14450 | 1 QLL94071. | 377 |
| CP051652.1:3239513-3240544 | **minus** | ABC transporter permease | **iolZ** | HER17_14455 | 1 QLL94072. | 343 |
| CP051652.1:3240557-3242104 | **minus** | sugar ABC transporter ATP-binding protein | **iolY** | HER17_14460 | 1 QLL94073. | 515 |
| CP051652.1:3242171-3243109 | **minus** | sugar ABC transporter substrate-binding protein | **iolX** | HER17_14465 | 1 QLL94074. | 312 |
| CP051652.1:3243190-3244176 | **minus** | inositol 2-dehydrogenase | **iolG** | HER17_14470 | 1 QLL94075. | 328 |
| CP051652.1:3245007-3246938 | **minus** | 3D-(3,5/4)-trihydroxycyclohexane-1,2-dione acylhydro | **iolD** | HER17_14475 | 1 QLL94076. | 643 |
| CP051652.1:3247041-3247301 | **minus** | hypothetical protein | | HER17_14480 | 1 QLL94077. | 86 |
| CP051652.1:3247415-3248926 | **minus** | CoA-acylating methylmalonate-semialdehyde dehydro | **iolA** | HER17_14485 | 1 QLL94078. | 503 |
| CP051652.1:3249254-3250111 | **plus** | MurR/RpiR family transcriptional regulator | **iolR** | HER17_14490 | 1 QLL94079. | 285 |
| CP051652.1:3250159-3250464 | plus | antibiotic biosynthesis monooxygenase | | HER17_14495 | 1 QLL94080. | 101 |
| CP051652.1:3250793-3251896 | plus | extracellular solute-binding protein | | HER17_14500 | 1 QLL94081. | 367 |
| CP051652.1:3252013-3253074 | minus | diguanylate cyclase AdrA | **adrA** | HER17_14505 | 1 QLL94082. | 353 |
| CP051652.1:3253308-3254144 | minus | PRD domain-containing protein | | HER17_14510 | 1 QLL94083. | 278 |
| CP051652.1:3254255-3255679 | minus | glycoside hydrolase family 1 protein | | HER17_14515 | 1 QLL94084. | 474 |
| CP051652.1:3255976-3257604 | minus | S8/S53 family peptidase | | HER17_14520 | 1 QLL94085. | 542 |
| CP051652.1:3257623-3258180 | minus | chorismate mutase | | HER17_14525 | 1 QLL94086. | 185 |
| CP051652.1:3258572-3260713 | minus | YjbH domain-containing protein | | HER17_14530 | 1 QLL94087. | 713 |
| CP051652.1:3260710-3261492 | minus | capsule biosynthesis GfcC family protein | | HER17_14535 | 1 QLL94088. | 260 |
| CP051652.1:3261502-3262179 | minus | YjbF family lipoprotein | | HER17_14540 | 1 QLL94089. | 225 |
| CP051652.1:3262255-3262548 | minus | hypothetical protein | | HER17_14545 | 1 QLL94090. | 97 |
| CP051652.1:3262997-3264403 | minus | NADP-dependent phosphogluconate dehydrogenase | **gndA** | HER17_14550 | 1 QLL94091. | 468 |

**Table S4.** ***Pectobacterium versatile iol*** **cluster genome location (tRNA-Asn), and addiction system (in green).**

| Genome location | Strand | Protein | Gene | Locus tag | Protein tag | amino acids |
|---|---|---|---|---|---|---|
| NZ_CP034276.1:1685041-1686447 | plus | NADP-dependent phosphogluconate dehydrogenase | **gndA** | EIP93_RS07610 | 1 WP_125232511.1 | 468 |
| NZ_CP034276.1:1686915-1687208 | plus | hypothetical protein | | EIP93_RS07615 | 1 WP_010296810.1 | 97 |
| NZ_CP034276.1:1687284-1687961 | plus | YjbF family lipoprotein | | EIP93_RS07620 | 1 WP_164496820.1 | 225 |
| NZ_CP034276.1:1687971-1688753 | plus | capsule biosynthesis GfcC family protein | | EIP93_RS07625 | 1 WP_119158778.1 | 260 |
| NZ_CP034276.1:1688750-1690873 | plus | YjbH domain-containing protein | | EIP93_RS07630 | 1 WP_125232512.1 | 707 |
| NZ_CP034276.1:1691266-1691823 | plus | chorismate mutase | | EIP93_RS07635 | 1 WP_040031358.1 | 185 |
| NZ_CP034276.1:1691843-1693471 | plus | S53 family peptidase | | EIP93_RS07640 | 1 WP_125232513.1 | 542 |
| NZ_CP034276.1:1693796-1695220 | plus | glycoside hydrolase family 1 protein | | EIP93_RS07645 | 1 WP_125232514.1 | 474 |
| NZ_CP034276.1:1695331-1696167 | plus | PRD domain-containing protein | | EIP93_RS07650 | 1 WP_125232515.1 | 278 |
| NZ_CP034276.1:1696401-1697462 | plus | diguanylate cyclase AdrA | **adrA** | EIP93_RS07655 | 1 WP_103860296.1 | 353 |
| NZ_CP034276.1:1697579-1698682 | minus | extracellular solute-binding protein | | EIP93_RS07660 | 1 WP_125232516.1 | 367 |
| NZ_CP034276.1:1699011-1699316 | minus | putative quinol monooxygenase | | EIP93_RS07665 | 1 WP_010296830.1 | 101 |
| NZ_CP034276.1:1699364-1700221 | minus | MurR/RpiR family transcriptional regulator | **iolR** | EIP93_RS07670 | 1 WP_010281425.1 | 285 |
| NZ_CP034276.1:1700549-1702060 | plus | CoA-acylating methylmalonate-semialdehyde dehydrogenase | **iolA** | EIP93_RS07675 | 1 WP_103860297.1 | 503 |
| NZ_CP034276.1:1702247-1704178 | plus | 3D-(3,5/4)-trihydroxycyclohexane-1,2-dione acylhydrolase | **iolD** | EIP93_RS07680 | 1 WP_103860298.1 | 643 |
| NZ_CP034276.1:1704885-1705871 | plus | inositol 2-dehydrogenase | **iolG** | EIP93_RS07685 | 1 WP_010308440.1 | 328 |
| NZ_CP034276.1:1705946-1706884 | plus | substrate-binding domain-containing protein | **iolX** | EIP93_RS07690 | 1 WP_010308444.1 | 312 |
| NZ_CP034276.1:1706957-1708504 | plus | sugar ABC transporter ATP-binding protein | **iolY** | EIP93_RS07695 | 1 WP_103972215.1 | 515 |
| NZ_CP034276.1:1708517-1709548 | plus | ABC transporter permease | **iolZ** | EIP93_RS07700 | 1 WP_010281439.1 | 343 |
| NZ_CP034276.1:1709589-1710722 | plus | **Gfo/Idh/MocA family oxidoreductase** | **idh** | EIP93_RS07705 | 1 WP_125232517.1 | 377 |
| NZ_CP034276.1:1710918-1712822 | plus | 5-dehydro-2-deoxygluconokinase | **iolC** | EIP93_RS07710 | 1 WP_039506680.1 | 634 |
| NZ_CP034276.1:1712982-1713872 | plus | myo-inosose-2 dehydratase | **iolE** | EIP93_RS07715 | 1 WP_125232518.1 | 296 |
| NZ_CP034276.1:1713960-1714796 | plus | 5-deoxy-glucuronate isomerase | **iolB** | EIP93_RS07720 | 1 WP_125232519.1 | 278 |
| NZ_CP034276.1:1714899-1715219 | minus | type II toxin-antitoxin system RelE/ParE family toxin | **parE** | EIP93_RS07725 | 1 WP_125232520.1 | 106 |
| NZ_CP034276.1:1715219-1715497 | minus | type II toxin-antitoxin system ParD family antitoxin | **parD** | EIP93_RS07730 | 1 WP_010304717.1 | 92 |
| NZ_CP034276.1:1715575-1716264 | minus | ABC transporter ATP-binding protein | | EIP93_RS07735 | 1 WP_103972220.1 | 229 |
| NZ_CP034276.1:1716286-1717983 | minus | iron ABC transporter permease | | EIP93_RS07740 | 1 WP_103941539.1 | 565 |
| NZ_CP034276.1:1717967-1719016 | minus | Fe(3+) ABC transporter substrate-binding protein | | EIP93_RS07745 | 1 WP_103941537.1 | 349 |
| NZ_CP034276.1:1719251-1720090 | minus | PRD domain-containing protein | | EIP93_RS07750 | 1 WP_103941535.1 | 279 |
| NZ_CP034276.1:1720185-1721648 | minus | family 1 glycosylhydrolase | | EIP93_RS07755 | 1 WP_103860310.1 | 487 |
| NZ_CP034276.1:1721648-1723510 | minus | beta-glucoside-specific PTS transporter subunit IIABC | | EIP93_RS07760 | 1 WP_125232521.1 | 620 |
| NZ_CP034276.1:1723951-1725015 | plus | sensor domain-containing diguanylate cyclase | | EIP93_RS07765 | 1 WP_125232522.1 | 354 |
| NZ_CP034276.1:1725201-1725539 | minus | DUF1869 domain-containing protein | | EIP93_RS07770 | 1 WP_125232523.1 | 112 |
| NZ_CP034276.1:1725550-1725882 | minus | DUF1971 domain-containing protein | | EIP93_RS07775 | 1 WP_010304744.1 | 110 |
| NZ_CP034276.1:1726264-1726854 | plus | flavin reductase family protein | | EIP93_RS07780 | 1 WP_125232524.1 | 196 |
| NZ_CP034276.1:1727090-1727656 | minus | TetR/AcrR family transcriptional regulator | | EIP93_RS07790 | 1 WP_116155736.1 | 188 |
| NZ_CP034276.1:1727794-1728885 | plus | alkene reductase | | EIP93_RS07795 | 1 WP_107332613.1 | 363 |
| NZ_CP034276.1:1728961-1729743 | plus | **SDR family oxidoreductase** | | EIP93_RS07800 | 1 WP_040031375.1 | 260 |
| NZ_CP034276.1:1729838-1730656 | plus | **oxidoreductase** | | EIP93_RS07805 | 1 WP_116585937.1 | 272 |
| NZ_CP034276.1:1730664-1731683 | plus | **NADP-dependent oxidoreductase** | | EIP93_RS07810 | 1 WP_116585936.1 | 339 |
| NZ_CP034276.1:1731956-1732031 | minus | **tRNA-Asn** | | EIP93_RS07815 | | |

#### Table S5. *Pectobacterium punjabense iol* cluster genome location (tRNA-Asn), and addiction system (in green).

| Genome location | Strand | Protein | Gene | Locus tag | Protein tag | amino acids |
|---|---|---|---|---|---|---|
| NZ_CP038498.1:1603971-1605377 | plus | NADP-dependent phosphogluconate dehydrogenase | gndA | E2566_RS07175 | 1 WP_107169886.1 | 468 |
| NZ_CP038498.1:1605836-1606111 | plus | hypothetical protein | | E2566_RS07180 | 1 WP_107169887.1 | 91 |
| NZ_CP038498.1:1606187-1606864 | plus | YjbF family lipoprotein | | E2566_RS07185 | 1 WP_107169888.1 | 225 |
| NZ_CP038498.1:1606874-1607656 | plus | capsule biosynthesis GfcC family protein | | E2566_RS07190 | 1 WP_107169889.1 | 260 |
| NZ_CP038498.1:1607653-1609776 | plus | YjbH domain-containing protein | | E2566_RS07195 | 1 WP_107169890.1 | 707 |
| NZ_CP038498.1:1610168-1610725 | plus | chorismate mutase | | E2566_RS07200 | 1 WP_107169891.1 | 185 |
| NZ_CP038498.1:1610746-1612362 | plus | S53 family peptidase | | E2566_RS07205 | 1 WP_107169892.1 | 538 |
| NZ_CP038498.1:1612658-1614082 | plus | glycoside hydrolase family 1 protein | | E2566_RS07210 | 1 WP_107169893.1 | 474 |
| NZ_CP038498.1:1614178-1615014 | plus | PRD domain-containing protein | | E2566_RS07215 | 1 WP_107169894.1 | 278 |
| NZ_CP038498.1:1615238-1616299 | plus | diguanylate cyclase AdrA | adrA | E2566_RS07220 | 1 WP_107169895.1 | 353 |
| NZ_CP038498.1:1616465-1617568 | minus | extracellular solute-binding protein | | E2566_RS07225 | 1 WP_107169896.1 | 367 |
| NZ_CP038498.1:1617895-1618200 | minus | putative quinol monooxygenase | | E2566_RS07230 | 1 WP_107169897.1 | 101 |
| NZ_CP038498.1:1618247-1619104 | minus | MurR/RpiR family transcriptional regulator | **iolR** | E2566_RS07235 | 1 WP_107169898.1 | 285 |
| NZ_CP038498.1:1619438-1620949 | plus | CoA-acylating methylmalonate-semialdehyde dehydrogenase | **iolA** | E2566_RS07240 | 1 WP_107169899.1 | 503 |
| NZ_CP038498.1:1621306-1623237 | plus | 3D-(3,5/4)-trihydroxycyclohexane-1,2-dione acylhydrolase (decyclizing) | **iolD** | E2566_RS07245 | 1 WP_107169900.1 | 643 |
| NZ_CP038498.1:1624031-1625017 | plus | inositol 2-dehydrogenase | **iolG** | E2566_RS07250 | 1 WP_107169947.1 | 328 |
| NZ_CP038498.1:1625098-1626036 | plus | substrate-binding domain-containing protein | **iolX** | E2566_RS07255 | 1 WP_107169901.1 | 312 |
| NZ_CP038498.1:1626109-1627656 | plus | sugar ABC transporter ATP-binding protein | **iolY** | E2566_RS07260 | 1 WP_107169902.1 | 515 |
| NZ_CP038498.1:1627669-1628700 | plus | ABC transporter permease | **iolZ** | E2566_RS07265 | 1 WP_107169903.1 | 343 |
| NZ_CP038498.1:1628740-1629873 | plus | Gfo/Idh/MocA family oxidoreductase | **idh** | E2566_RS07270 | 1 WP_107169904.1 | 377 |
| NZ_CP038498.1:1629913-1631817 | plus | 5-dehydro-2-deoxygluconokinase | **iolC** | E2566_RS07275 | 1 WP_107169905.1 | 634 |
| NZ_CP038498.1:1631853-1632743 | plus | myo-inosose-2 dehydratase | **iolE** | E2566_RS07280 | 1 WP_107169906.1 | 296 |
| NZ_CP038498.1:1632770-1633606 | plus | 5-deoxy-glucuronate isomerase | **iolB** | E2566_RS07285 | 1 WP_107169907.1 | 278 |
| NZ_CP038498.1:1633708-1634028 | minus | type II toxin-antitoxin system RelE/ParE family toxin | parE | E2566_RS07290 | 1 WP_107169908.1 | 106 |
| NZ_CP038498.1:1634028-1634306 | minus | type II toxin-antitoxin system ParD family antitoxin | parD | E2566_RS07295 | 1 WP_010281452.1 | 92 |
| NZ_CP038498.1:1634384-1635073 | minus | ABC transporter ATP-binding protein | | E2566_RS07300 | 1 WP_107169909.1 | 229 |
| NZ_CP038498.1:1635095-1636792 | minus | iron ABC transporter permease | | E2566_RS07305 | 1 WP_107169910.1 | 565 |
| NZ_CP038498.1:1636776-1637825 | minus | Fe(3+) ABC transporter substrate-binding protein | | E2566_RS07310 | 1 WP_107169911.1 | 349 |
| NZ_CP038498.1:1638059-1638898 | minus | PRD domain-containing protein | | E2566_RS07315 | 1 WP_107169948.1 | 279 |
| NZ_CP038498.1:1638963-1640426 | minus | family 1 glycosylhydrolase | | E2566_RS07320 | 1 WP_107169912.1 | 487 |
| NZ_CP038498.1:1640426-1642285 | minus | beta-glucoside-specific PTS transporter subunit IIABC | | E2566_RS07325 | 1 WP_107169913.1 | 619 |
| NZ_CP038498.1:1642739-1643785 | plus | sensor domain-containing diguanylate cyclase | | E2566_RS07330 | 1 WP_107169914.1 | 348 |
| NZ_CP038498.1:1643832-1644632 | minus | hypothetical protein | | E2566_RS07335 | 1 WP_107169915.1 | 266 |
| NZ_CP038498.1:1645058-1645396 | minus | DUF1869 domain-containing protein | | E2566_RS07340 | 1 WP_107169916.1 | 112 |
| NZ_CP038498.1:1645406-1645738 | minus | DUF1971 domain-containing protein | | E2566_RS07345 | 1 WP_107169917.1 | 110 |
| NZ_CP038498.1:1646119-1646709 | plus | flavin reductase family protein | | E2566_RS07350 | 1 WP_107169949.1 | 196 |
| NZ_CP038498.1:1646971-1647528 | plus | AAA family ATPase | | E2566_RS07355 | 1 WP_107169918.1 | 185 |
| NZ_CP038498.1:1647886-1648785 | minus | LysR family transcriptional regulator | | E2566_RS07360 | 1 WP_107169919.1 | 299 |
| NZ_CP038498.1:1649030-1649668 | plus | NAD(P)H-binding protein | | E2566_RS07365 | 1 WP_107169920.1 | 212 |
| NZ_CP038498.1:1649898-1650011 | plus | DUF1348 family protein | | E2566_RS21800 | 1 WP_240618647.1 | 37 |
| NZ_CP038498.1:1650183-1650258 | minus | **tRNA-Asn** | | E2566_RS07375 | | |

### Table S6. *Brenneria izbisi iol* cluster genome location (tRNA-Asn), associated mobile genetic element (integrase in red and T6SS) and addiction system (in green).

| Genome location | Strand | Protein | Gene | Locus tag | Protein tag | amino acids |
|---|---|---|---|---|---|---|
| NZ_JAMPJT010000002.1:288054-289460 | plus | NADP-dependent phosphogluconate dehydrogenase | gndA | NC803_RS04105 | 1 WP_264089113.1 | 468 |
| NZ_JAMPJT010000002.1:289985-290272 | plus | hypothetical protein | | NC803_RS04110 | 1 WP_264089112.1 | 95 |
| NZ_JAMPJT010000002.1:290348-291025 | plus | YjbF family lipoprotein | | NC803_RS04115 | 1 WP_264089111.1 | 225 |
| NZ_JAMPJT010000002.1:291035-291814 | plus | capsule biosynthesis GfcC family protein | | NC803_RS04120 | 1 WP_264089110.1 | 259 |
| NZ_JAMPJT010000002.1:291814-293943 | plus | YjbH domain-containing protein | | NC803_RS04125 | 1 WP_264089109.1 | 709 |
| NZ_JAMPJT010000002.1:294254-295093 | plus | PRD domain-containing protein | | NC803_RS04130 | 1 WP_264089108.1 | 279 |
| NZ_JAMPJT010000002.1:295224-296084 | minus | GHMP kinase | | NC803_RS04135 | 1 WP_264089107.1 | 286 |
| NZ_JAMPJT010000002.1:296077-297150 | minus | threonine-phosphate decarboxylase CobD | cobD | NC803_RS04140 | 1 WP_264089106.1 | 357 |
| NZ_JAMPJT010000002.1:297237-298301 | minus | NAD(P)-dependent alcohol dehydrogenase | | NC803_RS04145 | 1 WP_264089105.1 | 354 |
| NZ_JAMPJT010000002.1:298795-300231 | plus | cobyrinate a,c-diamide synthase | | NC803_RS04150 | 1 WP_264089104.1 | 478 |
| NZ_JAMPJT010000002.1:300228-301181 | plus | adenosylcobinamide-phosphate synthase CbiB | cbiB | NC803_RS04155 | 1 WP_264089103.1 | 317 |
| NZ_JAMPJT010000002.1:301196-301828 | plus | cobalt-precorrin-8 methylmutase | | NC803_RS04160 | 1 WP_264089102.1 | 210 |
| NZ_JAMPJT010000002.1:301825-302961 | plus | cobalt-precorrin-5B (C(1))-methyltransferase CbiD | cbiD | NC803_RS04165 | 1 WP_264089101.1 | 378 |
| NZ_JAMPJT010000002.1:302958-303560 | plus | cobalt-precorrin-7 (C(5))-methyltransferase | | NC803_RS04170 | 1 WP_264089100.1 | 200 |
| NZ_JAMPJT010000002.1:303550-304128 | plus | cobalt-precorrin-6B (C(15))-methyltransferase | | NC803_RS04175 | 1 WP_264089099.1 | 192 |
| NZ_JAMPJT010000002.1:304112-304921 | plus | cobalt-precorrin-4 methyltransferase | | NC803_RS04180 | 1 WP_264089098.1 | 269 |
| NZ_JAMPJT010000002.1:304902-305957 | plus | cobalt-precorrin 5A hydrolase | cbiG | NC803_RS04185 | 1 WP_264089097.1 | 351 |
| NZ_JAMPJT010000002.1:305957-306682 | plus | precorrin-3B C(17)-methyltransferase | | NC803_RS04190 | 1 WP_264089096.1 | 241 |
| NZ_JAMPJT010000002.1:306679-307473 | plus | cobalt-precorrin-6A reductase | | NC803_RS04195 | 1 WP_264089095.1 | 264 |
| NZ_JAMPJT010000002.1:307477-308271 | plus | sirohydrochlorin cobaltochelatase | | NC803_RS04200 | 1 WP_264089094.1 | 264 |
| NZ_JAMPJT010000002.1:308271-308990 | plus | cobalt-factor II C(20)-methyltransferase | | NC803_RS04205 | 1 WP_264089093.1 | 239 |
| NZ_JAMPJT010000002.1:308987-310504 | plus | cobyric acid synthase | | NC803_RS04210 | 1 WP_264089092.1 | 505 |
| NZ_JAMPJT010000002.1:310760-311302 | plus | bifunctional adenosylcobinamide kinase/adenosylcobina | cobU | NC803_RS04215 | 1 WP_264089091.1 | 180 |
| NZ_JAMPJT010000002.1:311299-312045 | plus | adenosylcobinamide-GDP ribazoletransferase | cobS | NC803_RS04220 | 1 WP_264089090.1 | 248 |
| NZ_JAMPJT010000002.1:312123-312731 | plus | adenosylcobalamin/alpha-ribazole phosphatase | | NC803_RS04225 | 1 WP_264089089.1 | 202 |
| NZ_JAMPJT010000002.1:312744-313808 | plus | nicotinate-nucleotide--dimethylbenzimidazole phosphorib | cobT | NC803_RS04230 | 1 WP_264089088.1 | 354 |
| NZ_JAMPJT010000002.1:314196-315059 | plus | MurR/RpiR family transcriptional regulator | | NC803_RS04235 | 1 WP_264089087.1 | 287 |
| NZ_JAMPJT010000002.1:315168-316301 | plus | ABC transporter substrate-binding protein | | NC803_RS04240 | 1 WP_264089086.1 | 377 |
| NZ_JAMPJT010000002.1:316301-317416 | plus | ABC transporter ATP-binding protein | | NC803_RS04245 | 1 WP_264089085.1 | 371 |
| NZ_JAMPJT010000002.1:317413-318261 | plus | ABC transporter permease | | NC803_RS04250 | 1 WP_264089084.1 | 282 |
| NZ_JAMPJT010000002.1:318258-319034 | plus | ABC transporter permease | | NC803_RS04255 | 1 WP_264089083.1 | 258 |
| NZ_JAMPJT010000002.1:319064-320209 | plus | Xaa-Pro peptidase family protein | | NC803_RS04260 | 1 WP_264089082.1 | 381 |
| NZ_JAMPJT010000002.1:320216-321076 | minus | MurR/RpiR family transcriptional regulator | iolR | NC803_RS04265 | 1 WP_264089081.1 | 286 |
| NZ_JAMPJT010000002.1:321408-322919 | plus | CoA-acylating methylmalonate-semialdehyde dehydroge | iolA | NC803_RS04270 | 1 WP_264089080.1 | 503 |
| NZ_JAMPJT010000002.1:322970-324901 | plus | 3D-(3,5/4)-trihydroxycyclohexane-1,2-dione acylhydrolas | iolD | NC803_RS04275 | 1 WP_264089079.1 | 643 |
| NZ_JAMPJT010000002.1:325443-326429 | plus | inositol 2-dehydrogenase | iolG | NC803_RS04280 | 1 WP_264089078.1 | 328 |
| NZ_JAMPJT010000002.1:326482-327438 | plus | substrate-binding domain-containing protein | iolX | NC803_RS04285 | 1 WP_264089077.1 | 318 |
| NZ_JAMPJT010000002.1:327627-329174 | plus | sugar ABC transporter ATP-binding protein | iolY | NC803_RS04290 | 1 WP_264089362.1 | 515 |
| NZ_JAMPJT010000002.1:329187-330218 | plus | ABC transporter permease | iolZ | NC803_RS04295 | 1 WP_264089076.1 | 343 |
| NZ_JAMPJT010000002.1:330234-331367 | plus | Gfo/Idh/MocA family oxidoreductase | idh | NC803_RS04300 | 1 WP_264089075.1 | 377 |
| NZ_JAMPJT010000002.1:331448-333352 | plus | 5-dehydro-2-deoxygluconokinase | iolC | NC803_RS04305 | 1 WP_264089074.1 | 634 |
| NZ_JAMPJT010000002.1:333390-334280 | plus | myo-inosose-2 dehydratase | iolE | NC803_RS04310 | 1 WP_264089073.1 | 296 |
| NZ_JAMPJT010000002.1:334390-335220 | plus | 5-deoxy-glucuronate isomerase | iolB | NC803_RS04315 | 1 WP_264089072.1 | 276 |
| NZ_JAMPJT010000002.1:335488-336363 | plus | MurR/RpiR family transcriptional regulator | | NC803_RS04320 | 1 WP_264089071.1 | 291 |
| NZ_JAMPJT010000002.1:336477-337055 | plus | D-alanyl-D-alanine dipeptidase | ddpX | NC803_RS04325 | 1 WP_264089361.1 | 192 |
| NZ_JAMPJT010000002.1:337140-338732 | plus | ABC transporter substrate-binding protein | | NC803_RS04330 | 1 WP_264089070.1 | 530 |
| NZ_JAMPJT010000002.1:338829-339845 | plus | ABC transporter permease | | NC803_RS04335 | 1 WP_264089069.1 | 338 |
| NZ_JAMPJT010000002.1:339851-340762 | plus | D,D-dipeptide ABC transporter permease | ddpC | NC803_RS04340 | 1 WP_264089068.1 | 303 |
| NZ_JAMPJT010000002.1:340764-341792 | plus | ABC transporter ATP-binding protein | | NC803_RS04345 | 1 WP_264089067.1 | 342 |
| NZ_JAMPJT010000002.1:341789-342730 | plus | ABC transporter ATP-binding protein | | NC803_RS04350 | 1 WP_264089066.1 | 313 |
| NZ_JAMPJT010000002.1:343675-344178 | plus | type VI secretion system contractile sheath small subuni | tssB | NC803_RS04355 | 1 WP_264089065.1 | 167 |
| NZ_JAMPJT010000002.1:344211-345689 | plus | type VI secretion system contractile sheath large subuni | tssC | NC803_RS04360 | 1 WP_264089064.1 | 492 |
| NZ_JAMPJT010000002.1:345695-346126 | plus | type VI secretion system baseplate subunit TssE | tssE | NC803_RS04365 | 1 WP_264089063.1 | 143 |
| NZ_JAMPJT010000002.1:346129-347895 | plus | type VI secretion system baseplate subunit TssF | tssF | NC803_RS04370 | 1 WP_264089062.1 | 588 |
| NZ_JAMPJT010000002.1:347859-348857 | plus | type VI secretion system baseplate subunit TssG | tssG | NC803_RS04375 | 1 WP_264089061.1 | 332 |
| NZ_JAMPJT010000002.1:348860-350077 | plus | type VI secretion system-associated FHA domain proteir | tagH | NC803_RS04380 | 1 WP_264089060.1 | 405 |
| NZ_JAMPJT010000002.1:350077-350598 | plus | type VI secretion system lipoprotein TssJ | tssJ | NC803_RS04385 | 1 WP_264089059.1 | 173 |
| NZ_JAMPJT010000002.1:350601-351941 | plus | type VI secretion system baseplate subunit TssK | tssK | NC803_RS04390 | 1 WP_264089058.1 | 446 |
| NZ_JAMPJT010000002.1:351954-352730 | plus | type IVB secretion system protein IcmH/DotU | icmH | NC803_RS04395 | 1 WP_264089057.1 | 258 |
| NZ_JAMPJT010000002.1:352745-355345 | plus | type VI secretion system ATPase TssH | tssH | NC803_RS04400 | 1 WP_264089056.1 | 866 |
| NZ_JAMPJT010000002.1:355348-356883 | plus | sigma 54-interacting transcriptional regulator | | NC803_RS04405 | 1 WP_264089055.1 | 511 |
| NZ_JAMPJT010000002.1:356883-357443 | plus | type VI secretion system-associated protein VasI | vasI | NC803_RS04410 | 1 WP_264089054.1 | 186 |
| NZ_JAMPJT010000002.1:357455-358873 | plus | type VI secretion system protein TssA | tssA | NC803_RS04415 | 1 WP_264089053.1 | 472 |
| NZ_JAMPJT010000002.1:358898-362395 | plus | type VI secretion system membrane subunit TssM | tssM | NC803_RS04420 | 1 WP_264089052.1 | 1165 |
| NZ_JAMPJT010000002.1:362436-363872 | plus | VasL domain-containing protein | | NC803_RS04425 | 1 WP_264089051.1 | 478 |
| NZ_JAMPJT010000002.1:363935-365296 | plus | hypothetical protein | | NC803_RS04430 | 1 WP_264089360.1 | 453 |
| NZ_JAMPJT010000002.1:365577-366095 | plus | Hcp family type VI secretion system effector | | NC803_RS04435 | 1 WP_264089050.1 | 172 |
| NZ_JAMPJT010000002.1:366257-368293 | plus | type VI secretion system tip protein VgrG | | NC803_RS04440 | 1 WP_264089049.1 | 678 |
| NZ_JAMPJT010000002.1:368303-369673 | plus | PAAR domain-containing protein | | NC803_RS04445 | 1 WP_264089048.1 | 456 |
| NZ_JAMPJT010000002.1:369673-370452 | plus | hypothetical protein | | NC803_RS04450 | 1 WP_264089047.1 | 259 |
| NZ_JAMPJT010000002.1:370521-371219 | plus | ankyrin repeat domain-containing protein | | NC803_RS04455 | 1 WP_264089046.1 | 232 |
| NZ_JAMPJT010000002.1:371524-372471 | plus | DUF4123 domain-containing protein | | NC803_RS04460 | 1 WP_264089045.1 | 315 |
| NZ_JAMPJT010000002.1:372468-373391 | plus | DUF4123 domain-containing protein | | NC803_RS04465 | 1 WP_264089044.1 | 307 |
| NZ_JAMPJT010000002.1:373404-378443 | plus | RHS repeat-associated core domain-containing protein | | NC803_RS04470 | 1 WP_264089043.1 | 1679 |
| NZ_JAMPJT010000002.1:378446-378775 | plus | hypothetical protein | | NC803_RS04475 | 1 WP_264089042.1 | 109 |
| NZ_JAMPJT010000002.1:378840-378974 | minus | uncharacterized gene | | NC803_RS04480 | | |
| NZ_JAMPJT010000002.1:379021-381225 | plus | uncharacterized gene | | NC803_RS04485 | | |
| NZ_JAMPJT010000002.1:381227-381595 | plus | barstar family protein | | NC803_RS04490 | 1 WP_264089041.1 | 122 |
| NZ_JAMPJT010000002.1:381670-381819 | minus | SymE family type I addiction module toxin | | NC803_RS17780 | 1 WP_318841717.1 | 49 |
| NZ_JAMPJT010000002.1:381924-382301 | plus | **tyrosine-type recombinase/integrase** | | NC803_RS04500 | 1 WP_264089040.1 | 125 |
| NZ_JAMPJT010000002.1:382722-383696 | plus | chemotaxis protein | | NC803_RS04505 | 1 WP_264089039.1 | 324 |
| NZ_JAMPJT010000002.1:384186-384593 | plus | hypothetical protein | | NC803_RS04510 | 1 WP_264089038.1 | 135 |
| NZ_JAMPJT010000002.1:384912-385826 | minus | LysR family transcriptional regulator | | NC803_RS04515 | 1 WP_264089037.1 | 304 |
| NZ_JAMPJT010000002.1:385931-386389 | plus | DMT family transporter | | NC803_RS04520 | 1 WP_264089036.1 | 152 |
| NZ_JAMPJT010000002.1:386394-386852 | plus | DMT family transporter | | NC803_RS04525 | 1 WP_264089035.1 | 152 |
| NZ_JAMPJT010000002.1:387411-387653 | plus | tautomerase family protein | | NC803_RS04530 | 1 WP_264089034.1 | 80 |
| NZ_JAMPJT010000002.1:387656-388597 | plus | pseudouridine-5'-phosphate glycosidase | | NC803_RS04535 | 1 WP_264089033.1 | 313 |
| NZ_JAMPJT010000002.1:388600-389286 | plus | HAD family hydrolase | | NC803_RS04540 | 1 WP_264089032.1 | 228 |
| NZ_JAMPJT010000002.1:389395-390612 | plus | questin oxidase family protein | | NC803_RS04545 | 1 WP_264089031.1 | 405 |
| NZ_JAMPJT010000002.1:390686-394567 | plus | amino acid adenylation domain-containing protein | | NC803_RS04550 | 1 WP_264089030.1 | 1293 |
| NZ_JAMPJT010000002.1:394800-395546 | minus | DNA-binding transcriptional regulator YciT | | NC803_RS04555 | 1 WP_264089029.1 | 248 |
| NZ_JAMPJT010000002.1:395618-396280 | minus | fructose-6-phosphate aldolase | fsa | NC803_RS04560 | 1 WP_264089028.1 | 220 |
| NZ_JAMPJT010000002.1:396425-397324 | plus | glycyl-radical enzyme activating protein | | NC803_RS04565 | 1 WP_264089027.1 | 299 |
| NZ_JAMPJT010000002.1:397329-399761 | plus | formate C-acetyltransferase/glycerol dehydratase family glycyl radical | | NC803_RS04570 | 1 WP_264089026.1 | 810 |
| NZ_JAMPJT010000002.1:400046-401509 | plus | family 1 glycosylhydrolase | | NC803_RS04575 | 1 WP_264089025.1 | 487 |
| NZ_JAMPJT010000002.1:401905-401980 | minus | **tRNA-Asn** | | NC803_RS04580 | | |
